# Optimal trade-off control in machine learning-based library design, with application to adeno-associated virus (AAV) for gene therapy

**DOI:** 10.1101/2021.11.02.467003

**Authors:** Danqing Zhu, David H. Brookes, Akosua Busia, Ana Carneiro, Clara Fannjiang, Galina Popova, David Shin, Kevin. C. Donohue, Edward F. Chang, Tomasz J. Nowakowski, Jennifer Listgarten, David. V. Schaffer

## Abstract

Adeno-associated viruses (AAVs) hold tremendous promise as delivery vectors for clinical gene therapy, but they need improvement. AAVs with enhanced properties, such as more efficient and/or cell-type specific infection, can be engineered by creating a large, diverse starting library and screening for desired phenotypes, in some cases iteratively. Although this approach has succeeded in numerous specific cases, such as infecting cell types from the brain to the lung, the starting libraries often contain a high proportion of variants unable to assemble or package their genomes, a general prerequisite for engineering any gene delivery goal. Herein, we develop and showcase a machine learning (ML)-based method for systematically designing more effective starting libraries — ones that have broadly good packaging capabilities while being as diverse as possible. Such carefully designed but general libraries stand to significantly increase the chance of success in engineering any property of interest. Furthermore, we use this approach to design a clinically-relevant AAV peptide insertion library that achieves 5-fold higher packaging fitness than the state-of-the-art library, with negligible reduction in diversity. We demonstrate the general utility of this designed library on a downstream task to which our approach was agnostic: infection of primary human brain tissue. The ML-designed library had approximately 10-fold more successful variants than the current state-of-the-art library. Not only should our new library serve useful for any number of other engineering goals, but our library design approach itself can also be applied to other types of libraries for AAV and beyond.

## Introduction

Adeno-associated viruses (AAVs) hold major promise as delivery vectors for gene therapy. While naturally-occurring AAVs can be clinically administered safely and in some cases efficaciously, they have a number of shortcomings that limit their use in many human therapeutic applications. For example, naturally-occurring AAVs do not target delivery to specific organs or cells, their delivery efficiency is limited, and they are susceptible to pre-existing neutralizing antibodies [1-3]. Consequently, directed evolution of the AAV capsid protein has emerged as a powerful strategy for engineering therapeutically suitable or optimal AAV variants. In directed evolution, a diversified library of AAV capsid sequences is subjected to multiple rounds of selection for a specific property of interest, with the aim of identifying and enriching the most effective variants [1, 4]. Primary techniques for constructing AAV starting libraries include error-prone PCR [1, 5], DNA shuffling [6, 7], structurally-guided recombination [8], peptide insertions [9], and phylogenetic reconstruction [10]. Recent studies have also explored computational strategies for setting the parameters that control the construction of these libraries. For example, genomic junctions that minimize AAV structure disruptions, suitable for recombination libraries, were computationally identified [9]. For mutagenesis libraries, genomic locations and their mutation probabilities were identified using single-substitution variant data, or by way of ancestral imputation from phylogenetic analysis [10-11].

Although successes have been achieved with directed evolution [4-6,8-9,13], several challenges are slowing progress [12]. For instance, a substantial fraction of the variants in the starting libraries for these selections are unable to assemble properly or package their payload efficiently — a basic requirement for any functional selection [11,14-15]. Consequently, much of the library is wasted, thereby decreasing the chance of successfully achieving any desired engineering goal in the downstream selections. Next Generation Sequencing (NGS) technologies enable analysis of properties for individual variants within a library, such as packaging fitness and infectivity, and the large quantity of data resulting from such assays suggest that machine learning (ML) could be a useful tool to help design more effective starting libraries for directed evolution. Herein, we propose a method to design such a ML-guided library that balances the requirements of packaging and diversity, to improve the probability of success in any general AAV directed evolution goal.

Recent studies have applied ML models trained on experimental data to generate novel AAV variants [16, 17]; however, these studies examined diversity post hoc, and provided no way to systematically navigate an optimal trade-off between diversity and packaging. In earlier work, Parker et al. [18] balanced “quality” and “novelty” in their library design. Quality was estimated from a statistical model evaluated on each sequence (specifically, a Potts model [19] trained on a fixed set of natural sequences), whereas novelty captured how different the library sequences were from naturally-occurring sequences but provided no indication of diversity within the library. Extensions of this work considered multiple fitness, or “quality,” scores [20]. In contrast to these, our approach will (i) allow for the use of any predictive model of fitness, (ii) explicitly address and control the diversity within the designed library, and (iii) be broadly applicable to different kinds of library construction.

We instantiated and evaluated our library design approach by designing a 7-mer peptide insertion library for AAV serotype 5 (AAV5) to optimally balance diversity and overall packaging fitness. Among the natural AAV serotypes, AAV5 has been suggested as a promising candidate for clinical gene delivery because of the low prevalence of pre-existing neutralizing antibodies and successful clinical development for hemophilia B [21-24]. We focus, specifically, on peptide insertion libraries because they are both simple and highly practical, having already been translated to the clinic (e.g., NCT03748784, NCT04645212, NCT04483440, NCT04517149, NCT04519749, NCT03326336, NCT05197270) [25].

Briefly our approach is as follows: first we assess the packaging fitness for variants in the current state-of-the art peptide insertion library class referred to as the NNK library. We then use these estimated packaging efficiencies as labels to build a predictive model from peptide insertion sequence to packaging fitness. Finally, we develop a design approach that can systematically trade off library diversity with packaging fitness, enabling us to choose an optimal trade-off. Our approach to ML-guided library design biases library construction towards variants that package well, thereby reducing the amount of wasted sequences and space in screening tasks. We show that our design approach yields a library with 5-fold higher packaging fitness than the NNK library, with negligible sacrifice to diversity, suggesting that our library will be more generally useful. As further evidence, when we subjected the NNK library to one round of packaging selection, the resulting pool of variants still had a lower packaging fitness than that of our initially designed library, while also being substantially less diverse. Finally, to demonstrate the general downstream utility of our designed library on an engineering task for which it was not designed, we showed in a primary human brain tissue selection, the ML-guided library yielded a 10-fold higher number of infectious variants compared to the NNK library, and these variants can be further selected for efficient and cell-specific infectivity. To the best of our knowledge, this is the first ML-guided AAV capsid library design used for selection in human tissue. While we focus on a therapeutically relevant capsid 7-mer peptide insertion library, our methods are general and can be applied to other AAV library types, and to proteins beyond AAV.

## Results

### AAV5-7mer peptide insertion library preparation and packaging selection

We used libraries with a variable 7-amino acid (7-mer) NNK sequence inserted at position 575-577 in the viral protein monomer, within a loop at the 3-fold symmetry axis associated with receptor binding and cell-specific entry [26, 27]. The “NNK” moniker refers to a broadly used strategy [28-30] involving a uniform distribution over all four nucleotides (N) in the first two positions of a codon, and equal probability on nucleotides G and T (K) in the third position; where the K in the third position was chosen to reduce the chance of stop codons which typically render the protein non-functional. The “NNK” moniker refers to a broadly used strategy [28-30] involving a uniform distribution over all four nucleotides (N) in the first two positions of a codon, and equal probability on nucleotides G and T (K) in the third position; where the K in the third position was chosen to reduce the chance of stop codons which typically render the protein non-functional. Each of the 7 amino acids in the insertion is sampled at random from this distribution during library construction. Although NNK libraries are among the most promising AAV libraries [2], a substantial fraction (>50%) of the variants in these libraries fail to package (i.e., do not assemble into viable capsids, and many more have lower packaging fitness than the parental virus [14, 15]. For example, placing a large hydrophobic residue in the 7-mer (solvent-exposed) region is likely destabilizing. Much of the experimental library is thus effectively wasted on poor fitness variants.

Our goal was to improve upon the commonly used NNK library and implicitly uncover a broad set of rules, as yet unknown, for insertion sequences that confer higher packaging fitness and then encode them in our library design so as to avoid such problems. In particular, our design approach will specify probabilities for each nucleotide in each position of the codon, at each position in the 7mer, in a manner that achieves better overall packaging than NNK, while maintaining high diversity. For example, we might specify for the first codon that the first nucleotide in the codon should be chosen with 20% chance as an A, 40% chance as a C, 35% chance as a T and 5% G; then specify four other such probabilities for the other two positions in the codon, for a total of 12 specified values. A designed library will specify these 84 () probabilities, which in turn will dictate the mean packaging fitness — through a complicated relationship that will be approximated with our machine learning predictive model—and library sequence diversity. We refer to designed libraries specified in this way as *position-wise nucleotide* specified.

First, we experimentally synthesized roughly 10^7^ variants from the NNK library to yield the NNK *pre-packaged library*. This plasmid library was then packaged, and the resulting viral particles were harvest ed and purified, and their genomes extracted, yielding the NNK *post-packaged library* (**Figure 1**) [31]. The sequences from both *pre-* and *post-*packaged libraries were then PCR amplified and deep sequenced (**Methods**).

**Figure 1:**
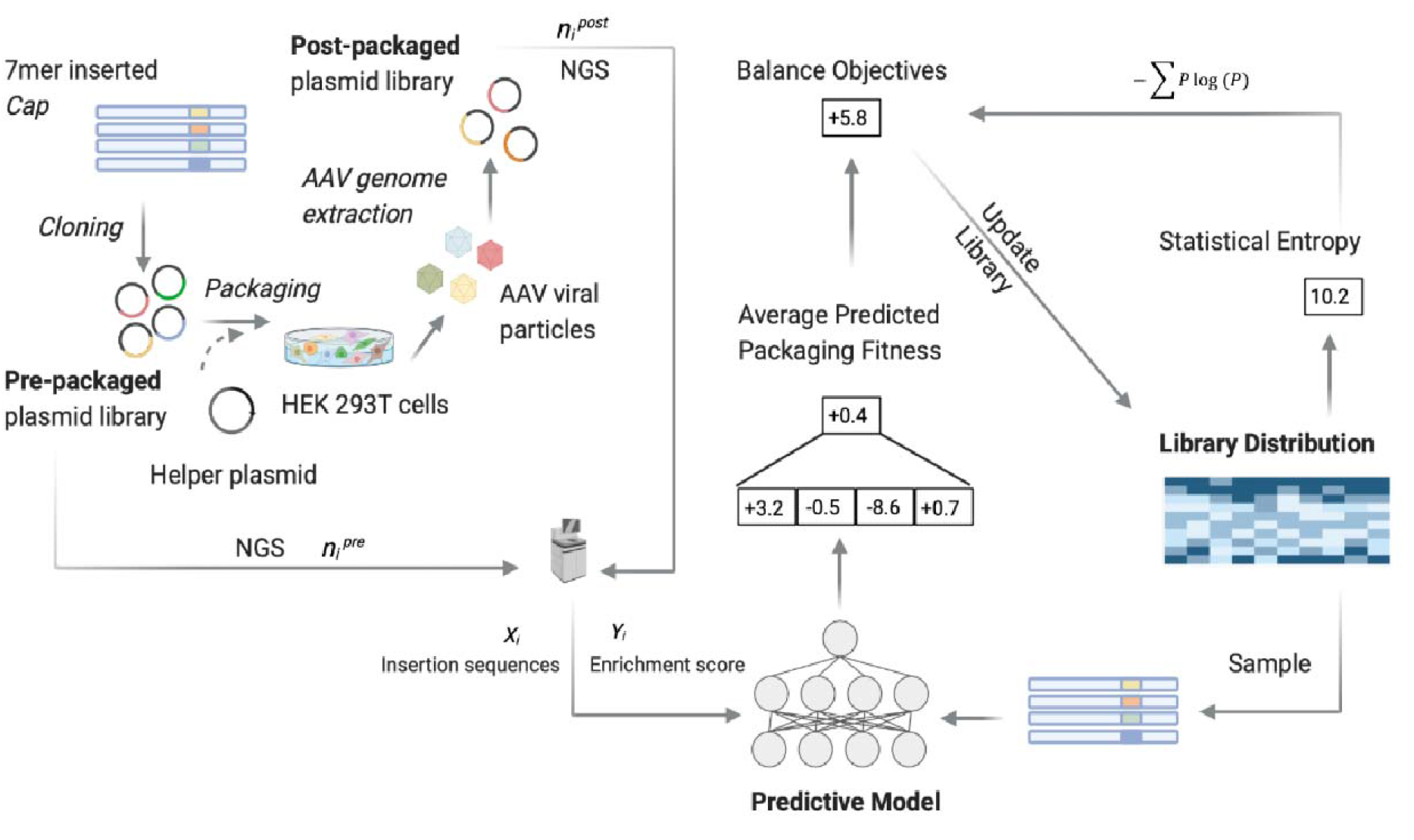
Experimental workflow for generating pre- and post-packaged AAV5-7mer library data for ML-based library design. NGS, next-generation sequencing; *N*_*i*_, number of reads for each unique insertion sequence *i*. Experimental data were used to build a supervised regression model where the target variable reflects the packaging success of each insertion sequence. The predictive model was then systematically inverted to design libraries that trace out an optimal trade-off curve between diversity and packaging fitness.

These experiments yielded 49,619,716 *pre-*packaged and 55,135,155 *post-*packaged sequencing reads, which collectively yielded read counts for 8,552,729 unique peptide sequences. For each unique sequence, we used the pre and post read counts to calculate a log enrichment score [11, 16, 32, 33] (**Methods**), a measure of its packaging fitness. Note, however, that a variant that appeared in 10 pre- and 100 post-packaged sequencing reads would have the same log enrichment score as one that appeared in 1 and 10 sequencing reads, even though the former has more data to support its value (*i*.*e*., is more stably statistically estimated). Consequently, we derived a procedure to take this into account in a statistically principled manner when estimating our regression model parameters. Our procedure assigns a weight to each unique sequence that is higher when the statistical estimate is more stable, and higher weighted sequences have more influence on the regression model (**Methods**). In the previous example, the variant with a read count ratio of 10:1 would get a smaller weight than the one with a ratio 100:10, as the former provides weaker evidence of enrichment.

### Training and evaluation of predictive model

To find the best model type to use for our ML-guided library design, we compared seven classes of ML regression models: three linear models and four feed-forward neural networks (NNs). Each model was trained using the log enrichment scores as the target variable and the sequence-specific weights described above. (**Methods**). The three linear models differed in the set of input features used. One used the “Independent Site” (IS) representation wherein individual amino acids in each 7-mer insertion sequence were one-hot encoded. Another used a “Neighbors” representation comprised of IS features, and additionally pairwise interactions between all positions that are directly adjacent in the amino acid sequence. The third used a “Pairwise” representation comprised of the IS features, and additionally all pairwise interactions among all positions in the sequence. All neural network models used the IS features alone, as these models have the capacity to construct higher-order interaction features from the IS features. Each NN architecture comprised exactly two densely connected hidden layers with tanh activation functions. The four NN models differed in the size of the hidden layers, with each using either 100, 200, 500, or 1000 nodes in both hidden layers.

We compared the performance of these seven models using the standard (unweighted) Pearson correlation between model predictions and true log enrichment scores on a held-out test set (training with weighted samples as described earlier). We randomly split the data into a training set containing 80% of the data points and a test set containing the remaining 20% of the points. Because our ultimate aim was to design a library of sequences that package well, we also studied how the models’ predictive accuracy changed when restricted to sequences in the test set with observed high packaging log enrichment. Specifically, we computed the Pearson correlation on subsets of the test set restricted to the fraction K of sequences with the highest observed log enrichment. By varying K, we traced out a performance curve where for lower K, the evaluation is more focused on accurate prediction of higher log enrichment scores rather than lower ones (**Figure 2a**). Overall, we found that the NN models performed better than the linear models, presumably owing to their capacity to construct more complex functions—in particular, to capture higher-order epistatic interactions in the fitness function. We selected “NN, 100” as our final model, as it performed similarly to the overall best-performing model, “NN, 1000”, but with many fewer parameters.

**Figure 2:**
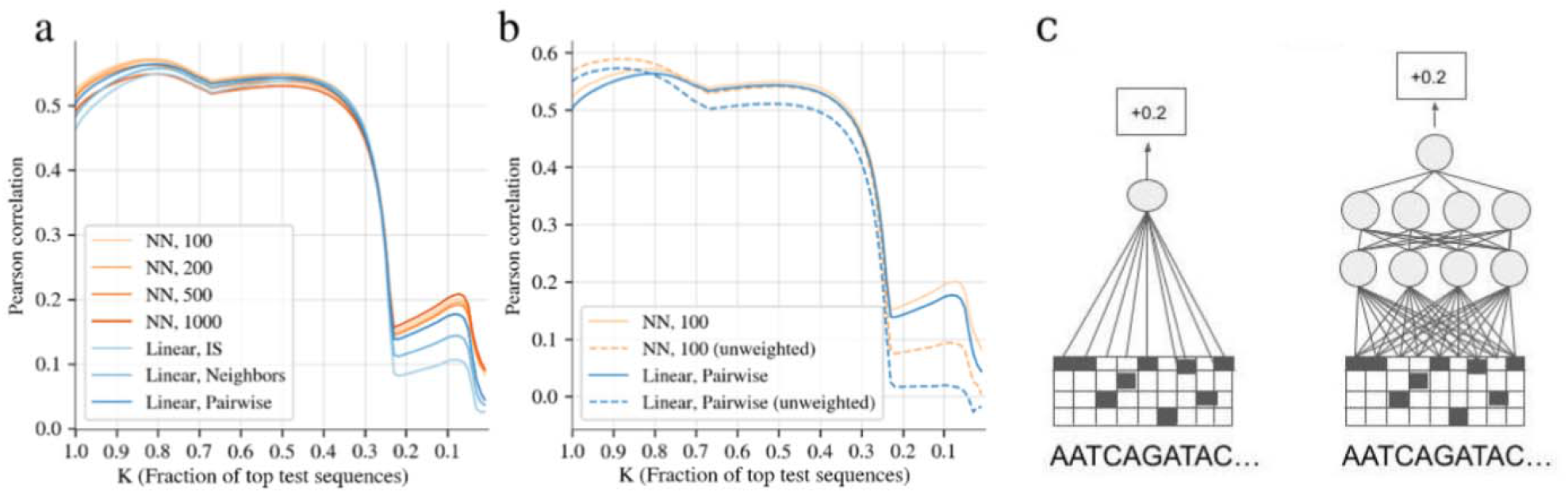
(a) Comparison of models for predicting AAV5-7mer packaging log enrichment scores, using Pearson correlation, and “top-K” Pearson correlation, where K, denotes what fraction of top-ranked observed log enrichment test sequences were used. The correlation is between predicted and true log enrichment scores. Seven different models including four neural network (NN) architectures, distinguished by the number of nodes in the hidden layers (100, 200, 500, 1000). (b) Similar plot to a) except comparing the use of weighted versus unweighted sequences during training, for the final selected model “NN, 100”, and a baseline “Linear, Pairwise”. (c) Schematic illustrations of the Linear, IS and NN, 4 predictive model.

Next we assessed the effect of training with our sequence-specific weights by retraining two of the models—the final model, “NN, 100” model and the “linear, Pairwise” model—this time with all weights set to 1.0 (i.e., unweighted), again using Pearson correlation to evaluate (**Figure 2b)**. Training in this unweighted manner, rather than weighted, resulted in a performance benefit for K near 1.0, but degraded the performance near K<0.25, a regime of particular interest since it focuses on variants with high log enrichment, and we ultimately aim to design a library that packages well (i.e., with high enrichment). These results further supported our choice of the weight trained “NN, 100” model with which to do library design.

### Experimental validation of the predictive models

Before proceeding to using our predictive model for library design, we first validated the “NN, 100” model by identifying and synthesizing five individual 7-mer insertion sequences that were not present in our experiment dataset. These five sequences were chosen to span a broad range of predicted log enrichment scores (−5.84 to 4.83 —see **Figure 3** for correspondence with viral titers). The five variants were packaged individually into viruses, harvested, and titered by quantifying the resulting number of genome-containing particles using digital-droplet PCR (**Methods**). High titer values indicated the variant was capable of packaging its genome properly in the assembled capsid. The agreement between model predictions and corresponding experimental measurement of vector titers (^4^ to ^11^ viral genomes (vg) /µL) (**Figure 3**) demonstrates that the predictive model was sufficiently accurate to be used for design.

**Figure 3:**
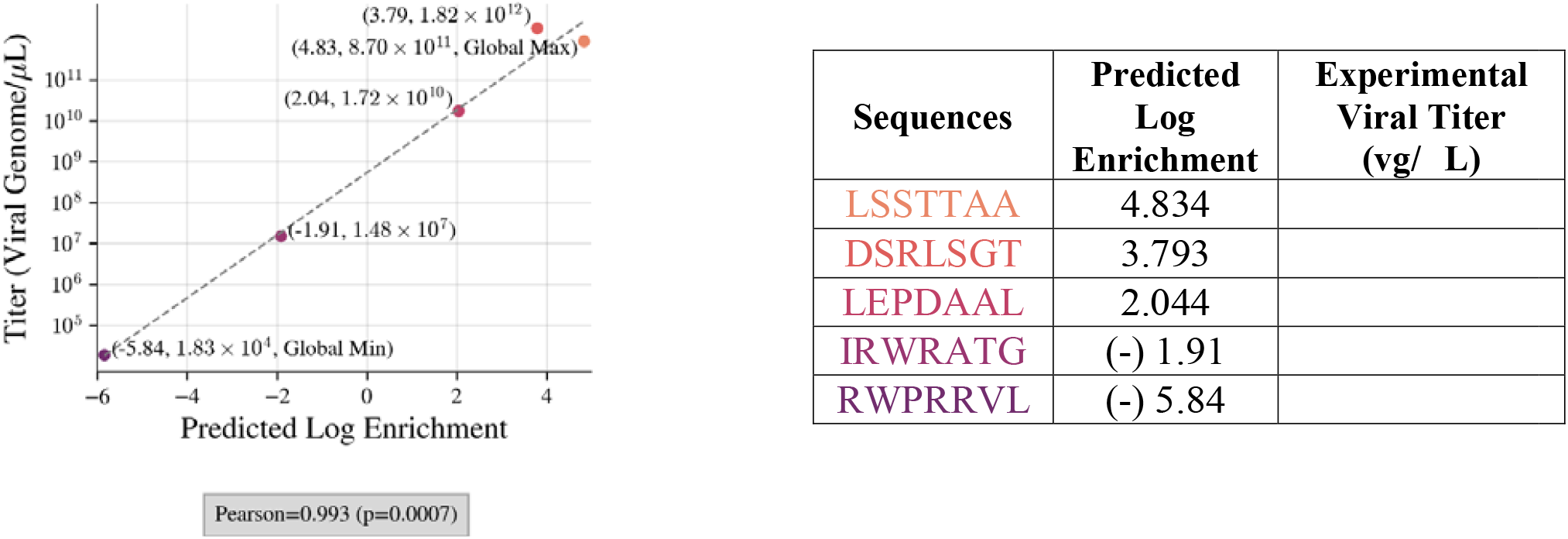
Experimental titers (vg /µL) versus predicted log enrichment scores for five variants selected to span a broad range of predicted log enrichment scores. Log enrichment scales are computed using natural logarithm.

### Model-guided library design

Having validated our final model for use with our library design task, we next aimed to design a library that packages better than the NNK library while maintaining good diversity. Inherent in this challenge is a trade-off between library diversity and mean predicted packaging fitness of the library. For example, note that mean predicted packaging fitness is maximized with a library that contains only a single variant with the highest predicted fitness, while diversity is maximized with a library uniformly distributed across sequence space, irrespective of packaging fitness. The library that is most effective for downstream selections will lie between these two extremes, balancing mean packaging fitnes with diversity. Because the best trade-off between these two extremes is not clear *a priori*, our approach to library design was to provide the tools to trace out an optimal trade-off curve, also known as a Pareto frontier (*e*.*g*., **Figure 4a**). Each point lying on this optimal frontier represents a library for which it is not possible to improve one desiderata (packaging or diversity), without hurting the other. Our Pareto optimal frontier, therefore, allows us to assess what mean library packaging fitness can be achieved for any given level of diversity. To generate each point (library) that lies on the optimal frontier curve, we define a library optimization objective that seeks to maximize mean predicted fitness subject to a library diversity constraint controlled by the value. This knob,, controls the trade off between library diversity and packaging ability; we set it to different values to trace out the Pareto frontier. We quantified the diversity of each theoretical library by computing the statistical entropy of the probabilistic distribution it corresponds to (**Methods**). We refer to this overall methodology enabling tracing out the optimal curve as *diversity-constrained optimal library design*. We note that the optimization problem is challenging to solve exactly (*i*.*e*., it is non-convex). Consequently, libraries computed as we trace out may not lie exactly on the optimal frontier. However, the frontier can nevertheless be inferred approximately, providing useful insights, as we shall see next.

**Figure 4:**
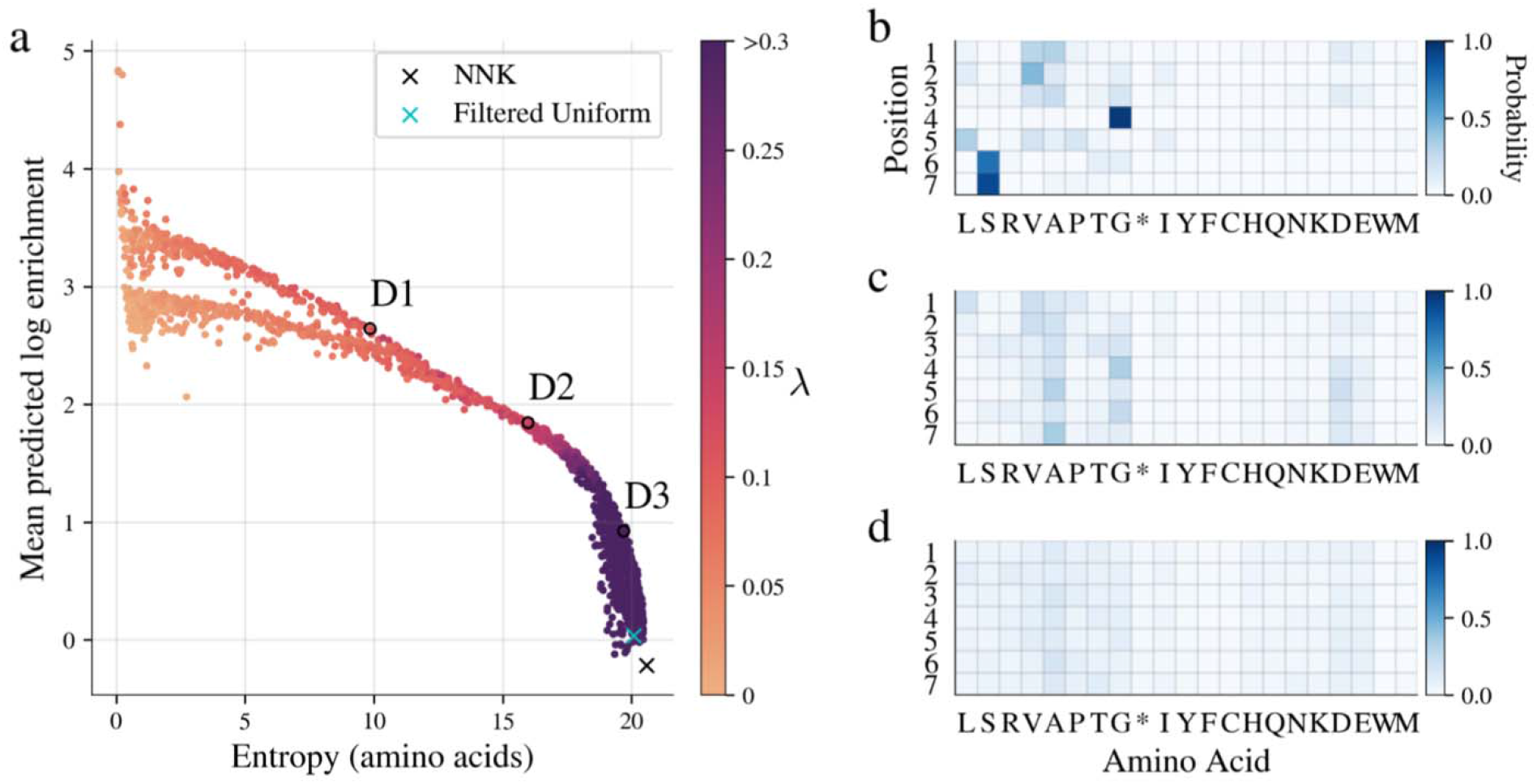
Designed AAV5-based 7-mer insertion libraries. Each point in a) represents a theoretical library designed with our diversity-constrained optimal library approach, with one particular diversity constraint, *λ*(higher values yields more diverse libraries). Entropy indicates diversity of the library distribution, while mean predicted log enrichment indicates overall library fitness; both quantities were computed from the theoretical library distribution. The baseline NNK library is denoted with a black “x”, while a cyan “x” denotes the “filtered uniform” library that is uniform over all 21-mer nucleotide sequences except for those containing stop codons. Three designed libraries have been circled and labeled D1-3 for reference. Due to the non-convex optimization problem, some dots are suboptimal (i.e., lie strictly below or to the left of other dots) and are therefore further from the optimal frontier, but are displayed for completeness. (b-d) designed library parameters (probability of each amino acid at each position) for the three designed libraries D1-3, respectively, highlighted in a).

We applied this diversity-constrained optimal library design methodology to the design of an improved AAV5-7mer peptide insertion library, yielding some striking implications (**Figure 4a**). We call out three designed libraries in particular — D1, D2, and D3 — as representative of three important areas of the curve and also show the NNK library overlayed. Remarkably, the NNK library has a dramatically poor mean predicted log enrichment (MPLE), much lower than any designed library. In contrast, library D3 had nearly identical diversity but substantially higher mean packaging fitness (top 50% of all designed libraries). This observation implies that D3 effectively dominates NNK in the sense that we increased the predicted packaging fitness without taking much loss to the diversity. Such concrete conclusions can be drawn from a Pareto frontier whenever one point on the frontier lies vertically above another. In addition, we see that compared to D3, D2 is less diverse but is predicted to package better (2.0-fold higher MPLE). Similarly, D1 is less diverse than D2, but again is predicted to package better (1.4-fold higher MPLE).

Although the original motivation for creating the NNK library was to reduce the number of stop codons, it does not eliminate them entirely. Therefore, for further comparison, we computed the mean packaging fitness and diversity of the theoretical library containing all possible sequences, except for any containing a stop codon. In practice, such a library is not physically realizable using this position-wide nucleotide specification strategy but serves as a useful comparator. We call this the “filtered uniform” library, and find that, on the one hand, it does have slightly higher mean packaging fitness than NKK, and correspondingly less diversity. However, these differences are negligible compared to the differences between NNK and D3, suggesting that further removal of stop codons is not the primary mechanism by which our ML-designed libraries achieve higher predicted packaging fitness.

### Richer library generation mechanisms

As mentioned earlier, each designed library specifies the 84 marginal probabilities of individual nucleotides at each position in the 21-bp insertion (**Table S2**). Our diversity-constrained optimal library design approach can, however, be used for any library construction method, such as one where we specify and synthesize individual 21-bp nucleotide sequences to create a library. We use the term “unconstrained” to refer to libraries that are designed with this construction method since individual synthesis offers the most control over sequences in the library. In contrast, a position-wise nucleotide specification strategy, such as the one we have used, cannot guarantee the inclusion of any particular sequence; we thus refer to libraries constructed in this manner as “constrained” libraries. We have focused our experiments on these constrained libraries because they are currently more cost-effective, and thus most widely used. Indeed, Weinstein *et al*. [34] showed that for a fixed cost, the use of a constrained library construction can yield orders of magnitude more promising leads in protein engineering than an unconstrained (individual synthesis) approach. As the cost of individual synthesis declines, it will become increasingly useful to use our design approach to specify unconstrained libraries that are both diverse and fit. With this future in mind, we also estimated the Pareto frontier for an unconstrained library (**Figure S5)**, which shows that, if cost were no concern, it would be advantageous to use an individual synthesis library construction approach, as its frontier substantially dominates that of our constrained library.

### Experimental validation of designed libraries

We synthesized two designed libraries (D2 and D3) from our optimality curve (**Figure 4a**) to assess the accuracy of the designed libraries’ trade-off between diversity and mean packaging fitness. Later we also test D2 in a downstream selection task of infecting brain tissue. The library D2 was chosen for being at the “elbow” of the curve, suggestive of a library making a good trade-off. The library D3 was chosen because, as discussed earlier, it dominates NKK by achieving much higher predicted packaging fitness with a negligible drop in diversity.

After experimentally constructing and deep sequencing these two designed libraries, we first checked that the physically realized library matched the statistics of the theoretical designed library distribution. Indeed, we found that the empirically observed position-wise probabilities for each amino acid in each of the designed libraries was within 5% of the designed specification (Table S3). Having validated that the constructed libraries were as specified, we packaged and harvested each library using the same methods as for the NNK library, yielding a pre- and post-packaged version of each. Next, we assessed to what degree the MPLE of each library reflected the measured library titers and found a strong positive Pearson correlation between them (r = 0.959, **Figure 5a**).

**Figure 5:**
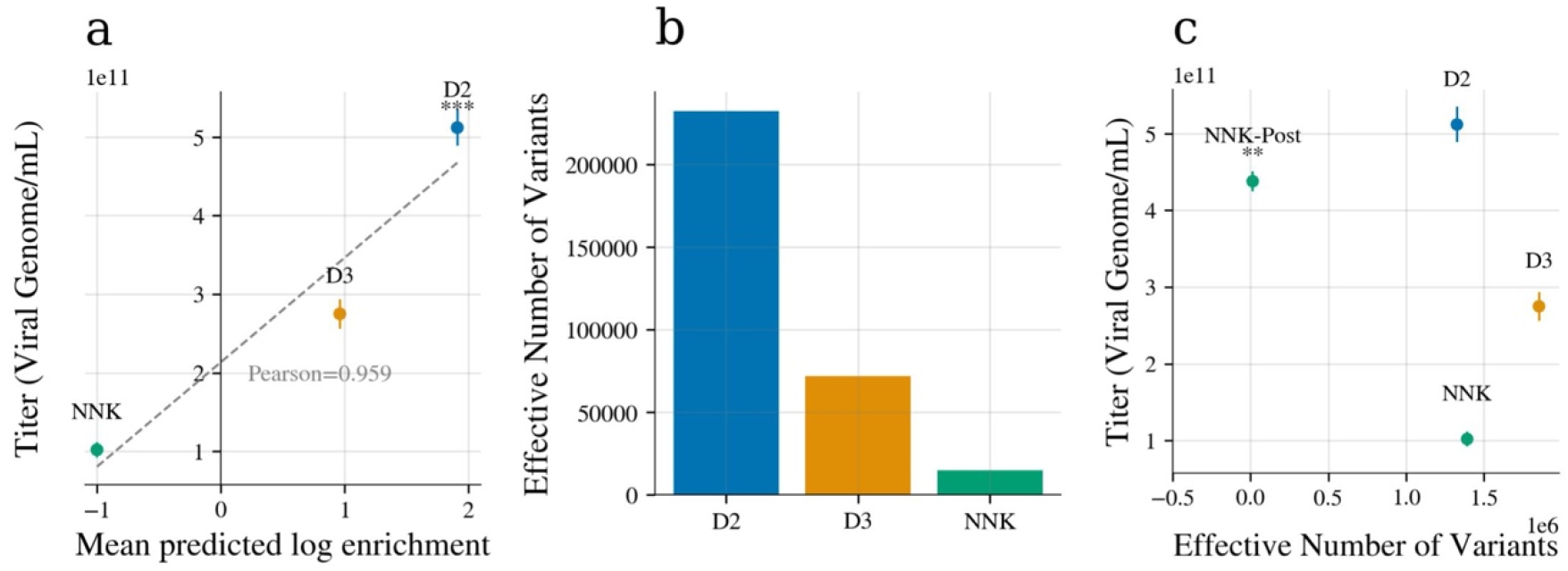
Comparison of ML-designed libraries D2 and D3 to the NNK library. (a) Experimental titers (viral genome/mL) plotted against the MPLE. (b) Comparison of the effective number of variants present in each library after packaging. (c) Experimental titers and effective number of variants for D2, D3, NNK, and NNK0post libraries. The NNK-post library represents the NNK library after one round of packaging selection. One-way ANOVA followed by Tukey test (** *p* <0.01 compared to D2, *** *p* <0.001 compared to NNK). In all cases, experimental titers are measured on 3 replicates. Graphs show means +/-SD.

As discussed earlier, D3 dominates the NNK library in fitness (one lies vertically above the other) and is thus predicted to be the better library. The choice between D3 and D2 is less clear, as they trade off packaging fitness and diversity. To assess such tradeoffs, we subjected each of D2, D3 and NNK to one round of packaging selection, and then estimated the effective number of variants remaining from the deep sequencing data (see **Methods**). A larger effective number of variants after selection suggests that a library contains more variants able to package.

Such an analysis revealed D2 to be better than D3 (**Figure 5b**). Consequently, we continued our comparison to NNK with only the D2 library.

Looking back at our measured titers, designed library D2 (MPLE∼ 2.0) showed a 5-fold higher packaging titer than that of the NNK library (MPLE∼ -0.9) with titers of 5.12 × 10 ^11^ and 1.02 × 10^11^ vg/mL, respectively. Next, we also measured the packaging titer of the NNK library after one round of packaging selection (NNK-post), finding that its titer (4.38 × 10 ^11^vg/mL) was lower than that of D2 (5.12 × 10^11^ vg/mL) (**Figure 5c**). This result suggests that the additional round of packaging was not enough to lift the NNK library’s titer level to that of library D2. Note, also, that the NNK-post library has only 1.48 × 10^4^ effective variants compared to the 1.33 × 10^6^ effective variants in D2. Collectively, these experimental results suggest that our ML-guided library design procedure yielded a more useful library than the NNK library, the peptide insertion library of choice for AAV directed evolution experiments.

### ML-designed AAV library for primary brain tissue infection

Having demonstrated our ability to design and construct libraries with better packaging and good diversity, we next investigated how these gains would translate into performance on a downstream selection task for which the library had not been tailored. After all, our goal was to design a generally useful library, agnostic to the downstream selection goal. Thus, we moved forward with two of the libraries, the NNK as a baseline and our designed library D2 and used each to infect primary adult brain tissue. Infecting such tissue with AAV can be a first step toward numerous clinical applications in the central nervous system. We applied each library onto human adult brain slices (**Figure S1, Methods)** and harvested the tissues after 72 hours of infection (**Figure 6a**). We evaluated the success of each library on this task by comparing the effective number of variants in each pool after infectivity selection. A higher effective number of variants post-brain infection would suggest that the starting library contained more variants that were able to successfully infect human brain tissue, indicating a more useful starting library and larger set of promising variants.

**Figure 6:**
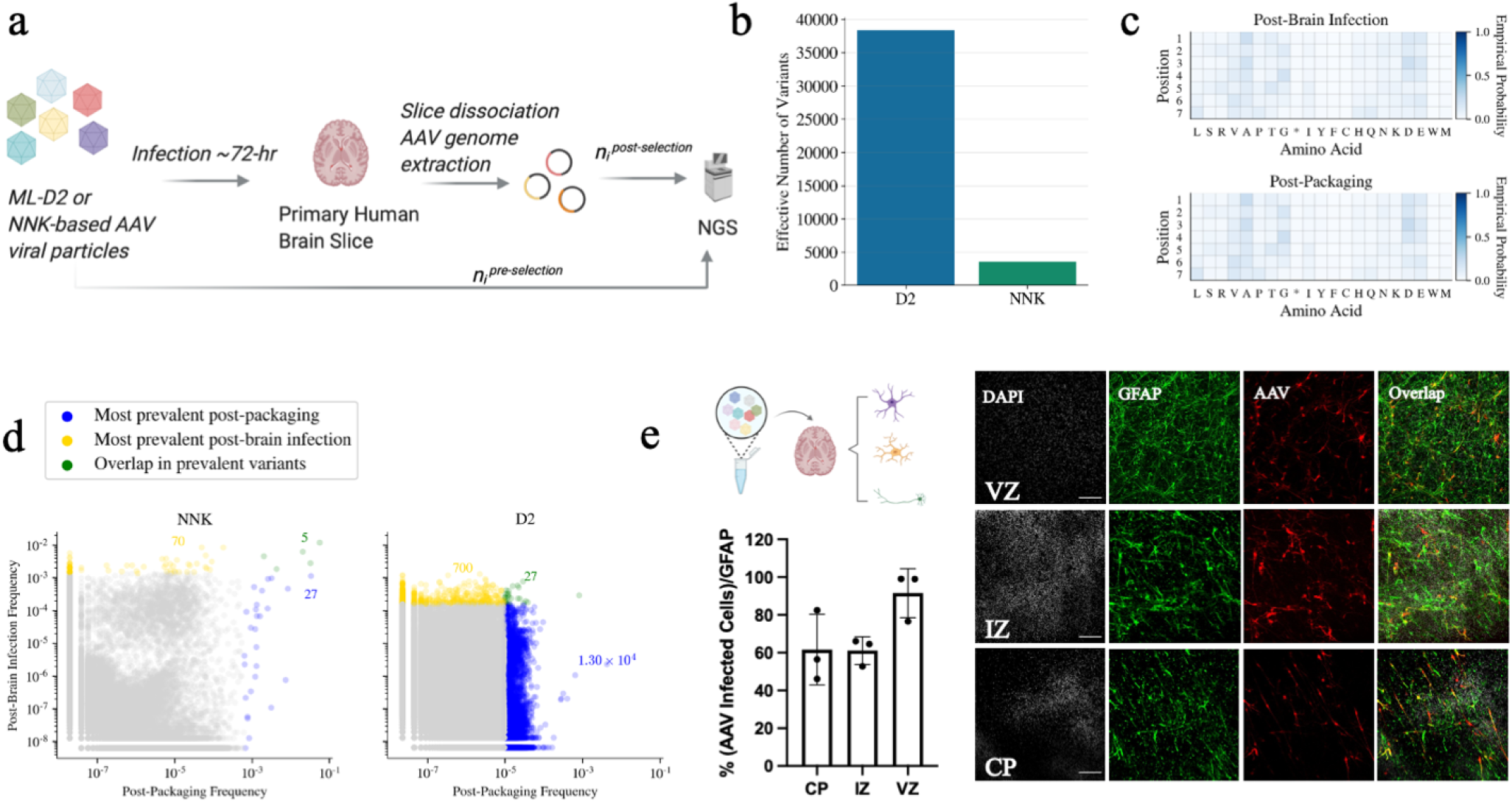
(a) General workflow of the primary adult brain infection study. (b) Effective number of variants (calculated from entropy) in NNK-post-brain infection vs. D2-post-brain infection; D2-post-brain infection exhibits a ∼10-fold increase in effective number of variants compared to that of NNK-post-brain infection. (c) Empirical probabilities of each amino acid at each position for D2 post-packaging and post-brain infection. (d) Scatterplots illustrating the behavior of individual variants over packaging and primary brain selection. Each axis shows the (log) prevalence of the variant in each library, as a fraction of reads in the library. For each library, variants in the top 20% are determined by first sorting unique variants by read count in descending order and then counting the number of unique variants comprising 20% of the total sequencing reads. Variants in the top 20% after packaging are colored blue, while those in the top 20% after brain selection are colored yellow. Those variants in the top 20% of both packaging and selection are colored green. The annotated colored numbers indicate the number of variants of each colored pool. A pseudocount of 1 was added to each variant in each library prior to plotting. See **Figure S4** for additional versions of (d) displaying variants in the top 50% and 80% of each library. (e) Cell-specific AAVs validation selected from the post-brain infection pool. (Green: Glial Fibrillary acidic protein (GFAP) marker; Red: AAV infected cells; Scale bar = 100 µm; CP: Cortical Plate; IZ: Intermediate Zone; VZ: Ventricular Zone).

We found that designed library D2 had a 10-fold higher post-brain infection effective number of variants than the NNK library (**Figure 6b, Figure S2)** — 38,350 vs. 3,541 effective variants. Diversity can be achieved in different ways, and we were interested to know whether diversity was spread out over the length of the 7-mer insertion, or if some positions might have “collapsed” to be more constrained as a result of the selection. Therefore, for each post-packaging and post-brain infection library, we computed the probability of each amino acid at each position and found largely uniform distributions over amino acids, thereby revealing that position-wise diversity was well-maintained (**Figure 6c, Figure S3**). We next compared the post-packaging and post-brain infection libraries at the level of individual variants to assess some practical implications of the difference in diversity between the NNK and D2 libraries (**Figure 6d**). We found a small set of variants dominated the post-packaging NNK library: the 32 most prevalent variants post-packaging (blue and green points in **Figure 6e**) accounted for 20% of the total sequencing reads. There were roughly 100-fold more unique variants in the top 20% of the D2 library post-packaging (1.32 × 10^4^ blue and green points), meaning that there is a much larger set of variants in the D2 library that package well when compared to the NNK library. In terms of downstream selection, the post-brain infection NNK library is dominated by a much smaller set of variants (∼10-fold fewer) compared to D2 (75 yellow and green points for NNK compared to 727 for D2). This suggests that the chances of discovering individual variants that successfully package and pass downstream selection are increased by using D2 instead of NNK as the starting library. In practice, then, the higher entropy for D2 post-packaging and post-brain infection translates to a much larger set of promising individual variants after each type of selection. We also considered the top 50% and 80% of the post-packaging and post-brain infection libraries (**Figure S4**) and found these conclusions to be consistent. Collectively, the results shown in **Figure 6** demonstrate that our designed library D2 provided more useful diversity over the widely used NNK library, thereby making it an effective, general starting library for downstream selections for which it was not specifically designed.

Finally, we validated that individual AAV variants from the ML-designed library D2 can not only package well but also successfully mediate cell-specific infection, which is a significant challenge in AAV engineering. For example, glial cells are important regulators of many aspects of human brain functions and diseases; however, true glial-cell specific targeting AAVs remain elusive [35]. To identify top variants for cell expression validation, we applied the D2 library to human brain tissue, dissociated and isolated glial cells, extracted the cells’ AAV genomes, and applied NGS. We ranked the variants in the D2-post-glia infection library by the enrichment score (computed between the initial D2 library and D2-post-glial infection library) and selected the top variants for individual validation. Each of these top selected glial specific AAV variants showed high titers (∼10× 10^12^ vg /µL) when packaged with a GFP-encoding genome (**Table S4**). Furthermore, immunostaining showed high levels of glial infection across multiple regions of the primary brain tissue (**Figure 6e**). Future work can extend our library design and selections to other cell types in brain or other tissues for a variety of therapeutic applications.

## Discussion

We developed a ML-based method for systematically designing diverse AAV libraries with good packaging capabilities, so that they can be used as starting libraries in directed evolution for engineering specific and enhanced AAV properties. A brief summary of our overall workflow was to (i) synthesize and sequence a baseline NNK library, the pre-packaged library; (ii) transfect the library into packaging cells (i.e., HEK 293T) to produce AAV viral vectors, harvest the successfully packaged capsids, extract viral genomes, and sequence to obtain the post-packaging library; (iii) build a supervised regression model where the target variable reflects the packaging success of each insertion sequence found; (iv) systematically invert the predictive model to design libraries that trace out an optimal trade-off curve between diversity and fitness; and (v) select a library design with a suitable tradeoff. We then validated both the predictive model and the designed library by experimentally measuring library packaging success and sequence diversity. Finally, we demonstrated that our ML-designed library is better able to infect primary human brain tissues as compared to the baseline NNK library.

In doing so, we have shown that (i) we can build accurate predictive models for AAV packaging fitness for 7mer insertion libraries; (ii) we can leverage these predictive models to design libraries that optimally trade off diversity with packaging fitness; and (iii) these designed libraries can be better starting libraries for downstream selection than standard libraries used today, despite not being tailored to the downstream task. To the best of our knowledge, this is the first work that develops and uses machine-learning based design to systematically identify a suite of optimal libraries along a trade-off curve of diversity and fitness. Additionally, it is the first to provide an end-to-end set of ML-based library design solutions, realized through experiments, in a therapeutically relevant system. We plan to generalize and apply this approach to further downstream selection tasks, including those relevant to gene replacement in the nervous system and evasion of pre-existing antibodies.

Our approach can, in principle, be used for other library construction techniques, such as individual gene sequence specification and synthesis (Figure S5). Our framework can also be extended to design libraries with multiple desired properties beyond diversity, by replacing the predictive model with one trained to simultaneously predict multiple functions or fitness combining such models when the properties are independent. This could be particularly useful to design libraries with improved cell sensitivity and specificity, which is particularly challenging using conventional experiment approaches.

## Acknowledgments

D.Z. was supported by Siebel Stem Cell Fellowship. D.Z., T.J.N. and J.L were supported by the Chan Zuckerberg Biohub. A.B., C.F., A.C. were supported by the National Science Foundation Graduate Research Fellowship Program under Grant No. DGE 1752814, Grant No. DGE 2146752, and Grant No. DGE 2146752; any opinions, findings, and conclusions or recommendations expressed in this material are those of the author(s) and do not necessarily reflect the views of the National Science Foundation. G.P. was supported by NIH NRSA F32 1F32MH118785. This work was supported, in part, by a grant from The Shurl and Kay Curci Foundation (T.J.N) and gifts from the Schmidt Futures and the William K. Bowes Jr Foundation (T.J.N.). Schematic illustrations were created with BioRender.com.

## Materials and Methods

### Construction of the NNK-based 7mer Insertion Library and Vector Packaging

We used libraries with a variable 7-amino acid (7-mer) insertion region flanked by amino acid linkers (TGGLS) introduced at position 575-577 in the viral protein monomer. (NNK)_7_ oligo was first synthesized (Elim) and introduced to the 5’ end of the right fragment by a primer overhang. Left and right fragments were each PCR amplified by primers Seq_F/Seq_R and 7mer_F/7mer_R, respectively (**Table S1**). PCR products of the two fragments were then purified individually and proceeded to the overlap extension PCR using Vent DNA polymerase (Thermofisher) with equimolar amounts of the left and right fragments for a total of 250ng DNA templates. The resulted library was then digested with *HindIII* and *NotI* and ligated into the replication competent AAV packaging plasmid pSub2. The resulting ligation reaction was electroporated into *Escherichia coli* for plasmid production and purification. Replication competent AAV was then packaged as been described previously [11, 26]. In short, AAV library vectors were produced by triple transient transfection of HEK293T cells with the addition of the pRepHelper, purified via iodixanol density centrifugation, and buffer exchanged into PBS by Amicon filtration.

### AAV Viral Genome Extraction and Titer

Packaged AAV vectors were first combined with equal volume of 10X DNase buffer (New England Biolabs, B0303S) and 0.5 µL 10 U/µL DNase I (New England Biolabs, M0303L) incubate for 30 min at 37 °C. Then equal volume of 2x Proteinase K Buffer was added with sample to break open capsid. After heat inactivating for 20 min at 95 °C, the sample was further diluted at 1:1000 and 1: 10,000 and use as templates for titer. DNase-resistant viral genomic titers were measured using digital-droplet PCR (ddPCR) (BioRad) using with Hex-ITR probes (CACTCCCTCTCTGCGCGCTCG) tagging the conserved regions of encapsidated viral genome of AAV. After primary tissue infection, capsid sequences were recovered by PCR from harvested cells using primers HindIII_F and NotI_R (**Table S1**). A ∼75-85 base pair region containing the 7mer insertion was PCR amplified from harvested DNA. Primers included the Illumina adapter sequences containing unique barcodes to allow for multiplexing of amplicons from multiple libraries. PCR amplicons were purified and sequenced with a single read run-on Illumina NovaSeq 6000.

### Data filtering and processing

The raw sequencing data consisted of 49,619,716 and 55,135,155 sequencing reads corresponding to the pre- and post-selection libraries, respectively. Each read contained (i) a 5 bp unique molecular identifier, (ii) a fixed 21 bp primer sequence, (iii) a 6 bp sequence representing the pre-insertion linker (two fixed amino acids that connect the insertion sequence to the capsid sequence at position 587), (iv) a variable 21 bp sequence containing the nucleotide insertion sequence, and (v) a 9 bp representing the post-insertion linker (three fixed amino acids that connect the insertion sequence to the capsid sequence at position 588). We filtered the reads, removing those that either contained more than 2 mismatches in the primer sequences or contained ambiguous nucleotides. After this filtering, the pre- and post-libraries contained 46,049,235 and 45,306,265 reads, respectively. The insertion sequences were then extracted from each read and translated to amino acid sequences.

### Log enrichment score and variance

We calculated the log enrichment scores (**Equation 1**) for each insertion sequence using the (filtered) sequencing data to quantify each sequence’s effect on packaging. Note that only 218,942 of the 8,552,729 unique sequences appear in both the pre- and post-selection libraries. A pseudo-count of 1 was added to each count so that the log enrichment score could still be calculated when the sequence appeared in only one of the libraries. In all cases, the natural log was used.

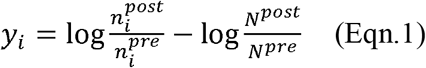

We estimated a variance associated with each log enrichment score using Equation 2, which follows by noting that each of the raw counts associated with a log enrichment score is a random variable. Specifically, the count associated with a sequence can be modeled as a Binomial random variable [32]. The log enrichment score (**Equation 1**) is then the log ratio of two Binomial random variables; it can be shown with the Delta Method [36] that, in the limit of infinite samples, the log ratio of two Binomial random variables converges in distribution to a Normal random variable with mean and variance approximated by Equations 1 and 2, respectively [32, 33].

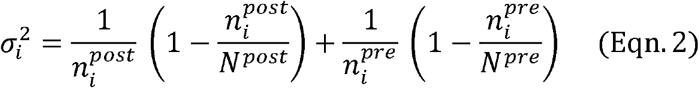

### Model training and evaluation

Our data processing yields a data set of the form 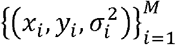 where the *x*_*i*_ are unique insertion sequences, *yi* are log enrichment scores associated with the insertion sequences, 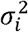 are the estimated variances of the log enrichment scores, and *M* = 8,555,729 is the number of unique insertion sequences in the data. We randomly split this data set into a training set containing 80% of the data and a test set containing the remaining 20% of the data.

We assume that the distribution of a log enrichment score given the associated insertion sequence is

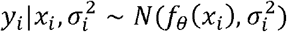

where *f*_*θ*_ is a function with parameters *θ* that parameterizes the mean of the distribution and represents a predictive model for log enrichment scores. We determined suitable settings of the parameters *θ* with Maximum Likelihood Estimation (MLE). The log-likelihood of the parameters of this model given the training set of *M* ′ ≤ *M* data points is given by

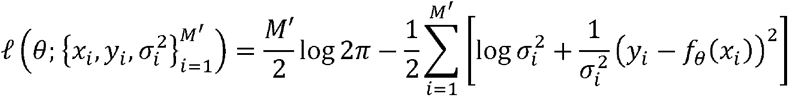

Performing MLE by optimizing this likelihood with respect to the model parameters, *θ*, results in the weighted least-squares loss function in **Equation 3**.

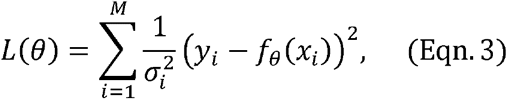

For the linear forms of *f*_*θ*_, the loss (**Equation 3**) is a convex function which can be solved exactly for the minimizing ML parameters. In order to stabilize training, we used a small amount of *l*_2_ regularization the Neighbors and Pairwise representations (with regularization coefficients 0.001 and 0.0025, respectively, chosen by cross-validation). For the neural network forms of *f*_*θ*_, the objective (**Equation 3**) is non-convex, and we use stochastic optimization techniques to solve for suitable parameters. We implemented these models in TensorFlow [37] and used the built-in implementation of the Adam algorithm [38] to approximately solve **Equation 3**.

To assess the prediction quality of each model, we calculated the Pearson correlation between the model predictions and observed log enrichment scores for different subsets of the sequences in the test set. Our aim is to use these models to design a library of sequences that package well (*i*.*e*., would be highly enriched in the post-selection library). We, therefore, assess how well the models perform for highly enriched sequences by progressively culling the test set to only include sequences with the largest observed log enrichment scores (**Figure 2**).

### Diversity-constrained optimal library design

We developed a general framework for sequence library design that (i) can be used with any predictive model of fitness, (ii) is broadly applicable to different library construction mechanisms (e.g., error prone PCR, site-specific marginal probability specification, individual synthesized sequences), and (iii) is simple to implement and extend. This framework balances mean predicted packaging fitness with entropy, a measure of diversity for probability distributions which has been used extensively in ecology to describe the diversity of populations [39]. Our approach is based on a maximum entropy formalism: we represent libraries as probability distributions and aim to find maximum entropy distributions that maximize entropy while also satisfying a constraint on the mean fitness, which is predicted by a user-specific model such as a neural network.

Let χ be the space of all sequences that may be included in a library (*e*.*g*., all amino acid sequences of length 7). We consider a library to be an abstract quantity represented by a probability distribution with support on χ. Let □ represent all such libraries and *p* ∈ 𝒫 one particular library. The entropy of this library is given by [40]:

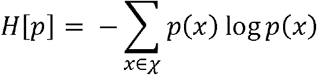

Now, let *f*(*x*)be a predictive model of fitness (*e*.*g*., from a trained neural network). Our goal is to find a diverse library, *p*, where the mean predicted fitness in the library, 𝔼 _*p*(*X*)_ [*f* (*x*)], is as high as possible. Formally, we want to find the library with the largest entropy such that the mean predicted fitness is above some cutoff. This objective is written

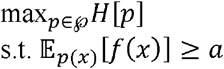

where *a* is the cutoff on the mean predicted fitness. It is straightforward to show that the solution to this optimization problem is given by [41]:

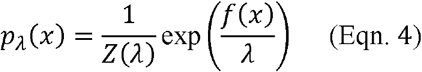

where *λ* > 0 is a Lagrange multiplier that is a monotonic function of the cutoff *a* and *Z* (*λ*) ∑_*x* ∈ χ_ exp (*f*(*x*)/*λ*) is a normalizing constant. **Equation 4** gives the probability mass of what is known as the maximum entropy distribution. The parameter *λ* controls the balance between diversity and mean fitness in the library (higher *λ* corresponds to more diversity). Each library, *p*_*λ*_, represents a point on a Pareto optimal frontier of libraries, which balances diversity and mean predicted fitness; these distributions cannot be perturbed in such a manner as to both increase the entropy and the mean fitness.

Theoretically, the entire Pareto frontier could be traced out by calculating the mean predicted fitness and entropy of *p* _*λ*_ for every possible setting of *λ*. In practice, we pick a discrete set of *λ* that traces out a practically useful curve.

As written so far, this framework can be used to select a particular library distribution, *p*_*λ*,_(*x*), with value *λ*, from the Pareto optimal curve. Then, if designing libraries comprised of individually specified sequences, one can sample individual sequences from this distribution, thereby designing a realizable, synthesizable library. However, for many cases of practical interest, it will not be cost-effective to synthesize individual sequences. We will, therefore, consider a more affordable library construction mechanism: a library of oligonucleotides is generated in a stochastic manner based on specified position-wise nucleotide probabilities. Because this position-wise nucleotide specification strategy does not allow one to specify individual sequences, we refer to libraries constructed in this way as *constrained*. In the next section, we describe how we use our design framework to set the parameters of these constrained libraries.

### Maximum Entropy Design for Constrained Libraries

In this section, we describe the design of libraries which are not specified at the level of individual sequences, but rather at the (less precise) level of position-specific distributions. In particular, we controlled the marginal probability of each nucleotide at each position. The probability mass function of the distribution representing a library specified by position-wise probabilities is given by:

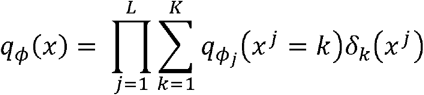

where L is the sequence length, K is the alphabet size (*i*.*e*., K = 4 for nucleotide libraries), *ϕ* ∈ ℝ ^*L* × *K*^ is a matrix of distribution parameters, *ϕ*_*j*_ is the j^th^ row of *ϕ, δ* _*k*_ (*x*^*j*^) = 1 if *x*^*j*^ = *k* and zero otherwise, and

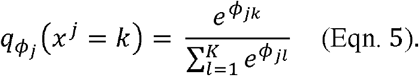

For an arbitrary predictive model (such as a neural network to predict log enrichment scores from sequence), the maximum entropy distribution (**Equation 4)** will generally not have the form of **Equation 5**. To apply the maximum entropy formulation to the design of libraries which are constrained to take a particular form, what we refer to as *constrained library* design, we take a variational approach: for a single, fixed value of *λ*, we find the constrained library distribution, *q*_*θ*_, that is the best approximation to the maximum entropy library distribution, *p*_*λ*_ in terms of the KL divergence,

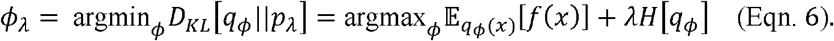

Our objective (**Equation 6**) is a non-convex function of the library parameters. The Stochastic Gradient Descent (SGD) algorithm has been shown to consistently find optimal or near-optimal solutions to a variety of non-convex problems, particularly in machine learning [42]. We use a variant of SGD based on the score function estimator [43] to solve **Equation 6**. We randomly initialize a parameter matrix, *ϕ* ^(0)^, with independent Normal samples, and then update the parameters according to

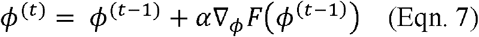

for *t* = 1, …, *T*, where we define 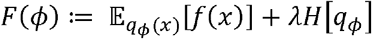 to be the objective function in **Equation 6**. The number of iterations, T, was set such that we observed convergence of the objective function values in most runs of the optimization. After *T* iterations, we assumed that we had reached a near-optimal solution (*i*.*e*., *ϕ* ^(*T*)^ can be used as an approximation of *ϕ*_*λ*_). The components of the gradient in **Equation 7** are given by

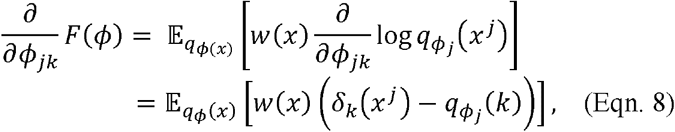

where we define the weights *w*(*x*) := *f*(*x*) − *λ* (1 + log*q*_*ϕ*_ (*x*))(**Supplementary Information**). The expectation in **Equation 8** cannot be solved exactly, so we use a Monte Carlo approximation:

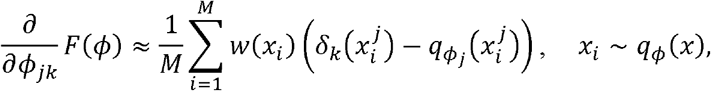

where *M* is the number of samples used for the MC approximation. We applied this maximum entropy framework to design site-specific marginal probability libraries of the 21 nucleotides corresponding to the 7 amino acid insertion using the (NN, 100) predictive model of fitness. **Figure 3** shows the near-optimal Pareto frontier resulting from 2,238 such library optimizations with *α* = 0.01, *T* = 2000, and *M* = 1000 and a range of settings of *λ*.

### Comparison of constructed libraries

Entropy is closely related to another notion of diversity known as *effective sample size*. The effective sample size of a library with entropy *H* is defined as *N*_*e*_ = *e*^*H*^, and corresponds to how many unique variants one would need to obtain entropy *H*, if each variant was constrained to have equal probability mass. This can be seen by noting that 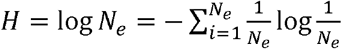.This interpretation of entropy is commonly used in the population genetics literature, first introduced by S Wright in 1931 [44].

When comparing designed theoretical libraries, we were able to compute the statistical entropy of each library distribution exactly in terms of its position-wise probabilities. However, when analyzing post-selection libraries, there is no known underlying probability distribution with which we can exactly compute entropy. Consequently, we instead estimated and compared the effective sample size of the empirically observed distribution in each library. Specifically, we estimated the effective number of samples in a library using the sequencing observations:

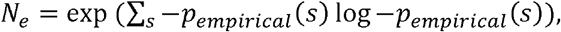

where *P* _*empirical*_ (*S*)corresponds to the empirical frequency of sequence *s* appearing in the post-selection sequencing data.

### Consent statement UCSF

De-identified tissue samples were collected with previous patient consent in strict observance of the legal and institutional ethical regulations. Sample use was approved by the Institutional Review Board at UCSF and experiments conform to the principles set out in the WMA Declaration of Helsinki and the Department of Health and Human Services Belmont Report.

### Primary Human Adult Brain Slices Culture and Library Infection

Adult surgical specimens from epilepsy cases were obtained from the UCSF medical center in collaboration with neurosurgeons with previous patient consent. Surgically excised specimens were immediately placed in a sterile container filled with N-methyl-D-glucamine (NMDG) substituted artificial cerebrospinal fluid (aCSF) of the following composition (in mM): 92 NMDG, 2.5 KCl, 1.25 NaH_2_PO_4_, 30 NaHCO_3_, 20 4-(2-hydroxyethyl)-1-piperazineethanesulfonic acid (HEPES), 25 glucose, 2 thiourea, 5 Na-ascorbate, 3 Na-pyruvate, 0.5 CaCl_2_·4H_2_O and 10 MgSO_4_·7H_2_O. The pH of the NMDG aCSF was titrated pH to 7.3–7.4 with 1M Tris-Base at pH8, and the osmolality was 300–305 mOsmoles/Kg. The solution was pre-chilled to 2–4□°C and thoroughly bubbled with carbogen (95% O_2_/5% CO_2_) gas prior to collection. The tissue was transported from the operating room to the laboratory for processing within 40–60□min. Blood vessels and meninges were removed from the cortical tissue, and then the tissue block was secured for cutting using superglue and sectioned perpendicular to the cortical plate to 300 μm using a Leica VT1200S vibrating blade microtome in aCSF. The slices were then transferred into a container of sterile-filtered NMDG aCSF that was pre-warmed to 32–34□°C and continuously bubbled with carbogen gas. After 12□min recovery incubation, slices were transferred to slice culture inserts (Millicell, PICM03050) on six-well culture plates (Corning) and cultured in adult brain slice culture medium containing 840 mg MEM Eagle medium with Hanks salts and 2mM L-glutamine (Sigma, M4642), 18 mg ascorbic acid (Sigma, A7506), 3 mL HEPES (1M stock) (Sigma, H3537), 1.68 mL NaHCO_3_ (892.75 mM solution, Gibco, 25080-094), 1.126 mL D-glucose, (1.11M solution, Gibco, A24940-01), 0.5 mL penicillin/streptomycin, 0.25 mL GlutaMax (at 400x, Gibco, 35050-061), 100 μL 2M stock MgSO_4_.7H_2_O (Sigma, M1880), 50 μL 2M stock CaCl_2_.2H_2_O (Sigma, C7902), 50 μL insulin from bovine pancreas, (10 mg/mL, Sigma, I0516), 20 mL horse serum-heat inactivated, 95 mL MilliQ H_2_O (as previously described [45]). The following day after plating, adult human brain slices were infected with the viral library at an estimated of 10,000 MOI (N=3 per group) based on the number of cells estimated per slice. Slices were cultured at the liquid–air interface created by the cell-culture insert in a 37°C incubator at 5% CO_2_ for 72 hours post infection.

### Slice Culture Dissociation, Cell Purification and Hirt Extraction

Seventy-two hours after infection with the viral library, cultured brain tissue slices were first rinsed with DPBS (Gibco, 14190250) twice and detached from the filters. Then mechanically minced to 1mm^2^ pieces and enzymatically digested with papain digestion kit (Worthington, LK003163) with the addition of DNase for 1 hr at 37°C. After the enzymatic digestion, tissue was mechanically triturated using fire-polished glass pipettes (Fisher Scientific, cat#13-678-6A), filtered through a 40 μm cell strainer (Corning 352340), pelleted at 300xg for 5 minutes and washed twice with DBPS. Following mechanical digestion, the slices were first treated with lysis buffer (10% SDS, 1M Tris-HCL, pH 7.4-8.0, and 0.5M EDTA, pH 8.0) with the addition of RNase A (Thermo Scientific, EN0531) for 60 min at 37 °C and proteinase K (New England Biolabs, P8107S) for 3 hours at 55 °C. The enzymatically digested tissue homogenate was then proceeded to the Hirt column protocol as previously published [46].

### Primary prenatal brain slices

Deidentified primary tissue samples were collected with previous patient consent in strict observance of the legal and institutional ethical regulations. Cortical brain tissue was immediately placed in a sterile conical tube filled with oxygenated artificial cerebrospinal fluid (aCSF) containing 125 mM NaCl, 2.5 mM KCl, 1mM MgCl_2_, 1 mM CaCl_2_, and 1.25 mM NaH_2_PO4 bubbled with carbogen (95% O_2_/5% CO_2_). Blood vessels and meninges were removed from the cortical tissue, and then the tissue block was embedded in 3.5% low-melting-point agarose (Thermo Fisher, BP165-25) and sectioned perpendicular to the ventricle to 300 μm using a Leica VT1200S vibrating blade microtome in a sucrose protective aCSF containing 185 mM sucrose, 2.5 mM KCl, 1 mM MgCl_2_, 2 mM CaCl_2_, 1.25 mM NaH_2_PO_4_, 25 mM NaHCO_3_, 25 mM d-(+)-glucose. Slices were transferred to slice culture inserts (Millicell, PICM03050) on six-well culture plates (Corning) and cultured in prenatal brain slice culture medium containing 66% (vol/vol) Eagle’s basal medium, 25% (vol/vol) HBSS, 2% (vol/vol) B27, 1% N2 supplement, 1% penicillin/streptomycin and GlutaMax (Thermo Fisher). Slices were cultured in a 37 °C incubator at 5% CO_2_, 8% O_2_ at the liquid–air interface created by the cell-culture insert.

### Slice dissociation and cell purification

Cultured brain slices were washed twice with DPBS (Gibco, 14190250), detached from the filters and enzymatically digested with papain digestion kit (Worthington, LK003163) with the addition of DNase for 30 mins at 37°C. Following enzymatic digestion, slices were mechanically triturated using a fire-polished glass pipette, filtered through a 40 μm cell strainer test tube (Corning 352235), pelleted at 300xg for 5 minutes and washed twice with DBPS.

Dissociated cells were resuspended in MACS buffer (DPBS with 1 mM EGTA and 0.5% BSA) with addition of DNAse and incubated with CD11b antibody (microglia) for 15 minutes on ice. After the incubation, cells were washed in a 10 ml of MACS buffer and loaded on LS columns (Miltenyi Biotec, 130-042-401) on the magnetic stand. Cells were washed 3 times with 3 ml of MACS buffer, then the column was removed from the magnetic field and microglia cells were eluted using 5 ml of MACS buffer. The flow-through cells were then gently prepared to separate out neurons using polysialylated-neural cell adhesion molecule (PSA-NCAM), and the flow-through cell population was used as glial-cell type. Cells were pelleted, re-suspended in 1 ml of culture media and counted.

### Immunofluorescence and Antibodies

Primary human brain slices were fixed on the filters in 4% PFA for 1 hour at room temperature and washed 3x with PBS for 5 mins each wash. Slices were carefully detached from the culture filter inserts and places into 12 well plates. Blocking and permeabilization were performed in a blocking solution consisting of 10% normal donkey serum, 1% Triton X-100, and 0.2% gelatin for 1 hour. Primary and secondary antibodies were diluted and incubated in the blocking solution. Prenatal brain slices were incubated with primary antibodies at 4^0^C overnight, washed 3x with washing buffer (1% Triton X-100 in PBS). Adult brain slices were incubated with primary antibodies for two days and washed 3x with washing buffer (1% Triton X-100 in PBS). Slices were incubated with secondary antibodies in the blocking buffer at 4^0^C overnight and washed with washing buffer for 5x for 10 mins each. Images were collected using Leica SP8 confocal system with 10x and 20x air objective and processed using ImageJ/Fiji and Affinity Designer software. Primary antibodies used in this study included: chicken GFAP (1:1,000, Abcam, ab4674), rabbit dsRed (1:250, Takara, 632496), and DAPI. Secondary antibodies were species-specific AlexaFluor secondary antibodies (1:2,000, ThermoFisher).

## Supplementary Materials

**Table S1.**
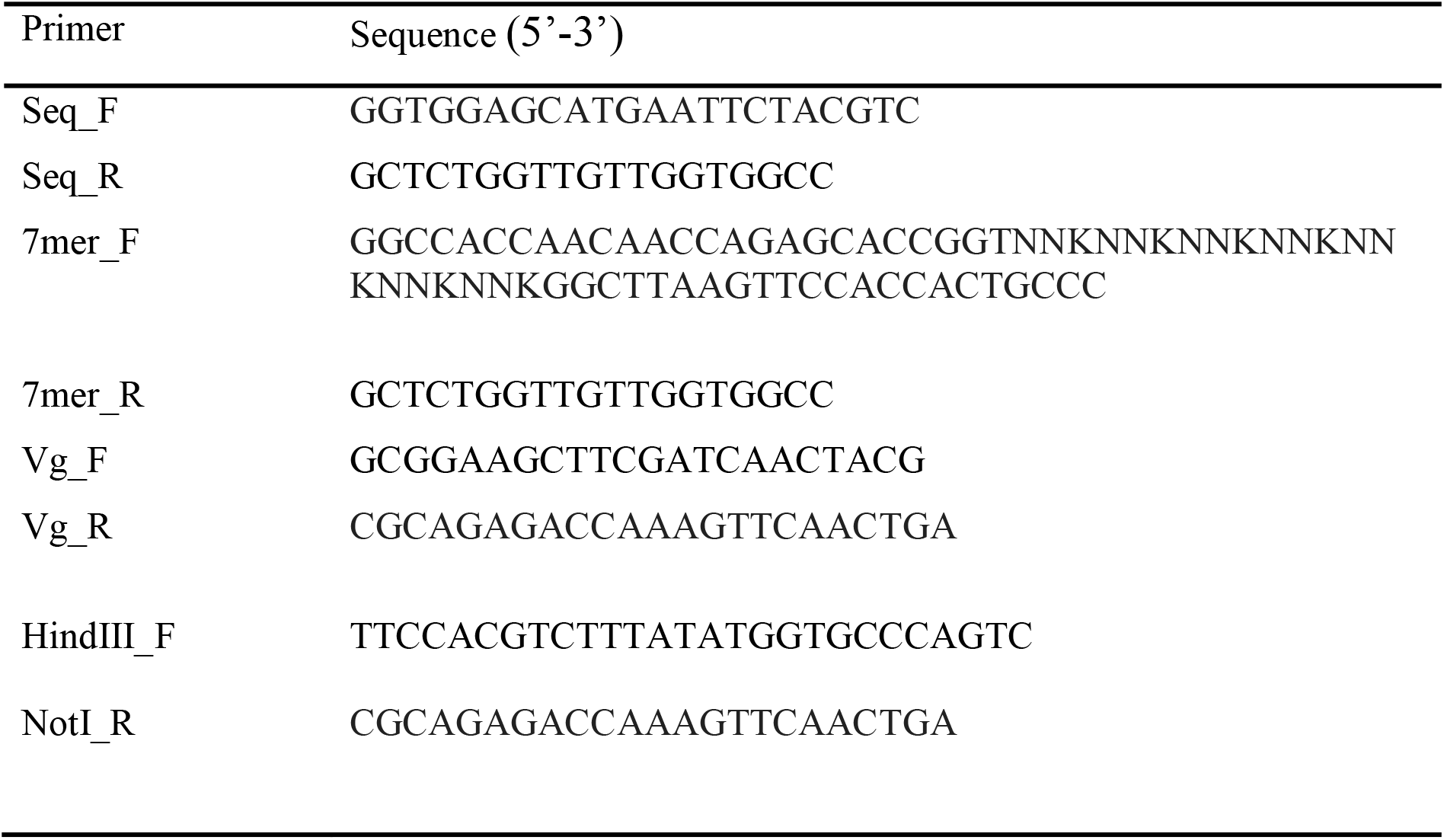
Primer sequences for PCR reactions. Primer Sequence (5’-3’)

**Table S2.**
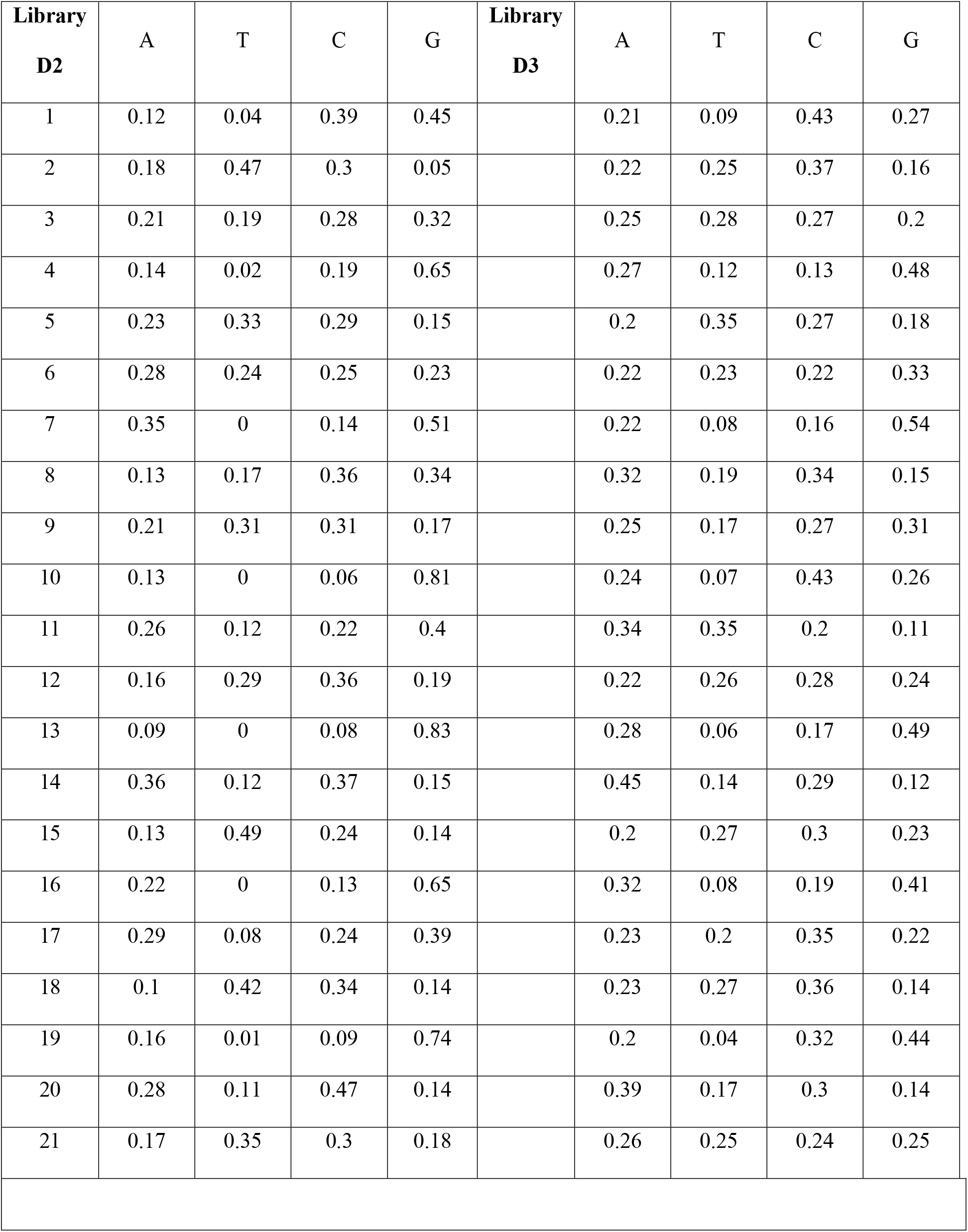
Marginal probabilities of library D2 and D3 nucleotides at 21-bp position chart.

| **Library**<br>**D2** | A | T | C | G | **Library**<br>**D3** | A | T | C | G |
| --- | --- | --- | --- | --- | --- | --- | --- | --- | --- |
| 1 | 0.12 | 0.04 | 0.39 | 0.45 |  | 0.21 | 0.09 | 0.43 | 0.27 |
| 2 | 0.18 | 0.47 | 0.3 | 0.05 |  | 0.22 | 0.25 | 0.37 | 0.16 |
| 3 | 0.21 | 0.19 | 0.28 | 0.32 |  | 0.25 | 0.28 | 0.27 | 0.2 |
| 4 | 0.14 | 0.02 | 0.19 | 0.65 |  | 0.27 | 0.12 | 0.13 | 0.48 |
| 5 | 0.23 | 0.33 | 0.29 | 0.15 |  | 0.2 | 0.35 | 0.27 | 0.18 |
| 6 | 0.28 | 0.24 | 0.25 | 0.23 |  | 0.22 | 0.23 | 0.22 | 0.33 |
| 7 | 0.35 | 0 | 0.14 | 0.51 |  | 0.22 | 0.08 | 0.16 | 0.54 |
| 8 | 0.13 | 0.17 | 0.36 | 0.34 |  | 0.32 | 0.19 | 0.34 | 0.15 |
| 9 | 0.21 | 0.31 | 0.31 | 0.17 |  | 0.25 | 0.17 | 0.27 | 0.31 |
| 10 | 0.13 | 0 | 0.06 | 0.81 |  | 0.24 | 0.07 | 0.43 | 0.26 |
| 11 | 0.26 | 0.12 | 0.22 | 0.4 |  | 0.34 | 0.35 | 0.2 | 0.11 |
| 12 | 0.16 | 0.29 | 0.36 | 0.19 |  | 0.22 | 0.26 | 0.28 | 0.24 |
| 13 | 0.09 | 0 | 0.08 | 0.83 |  | 0.28 | 0.06 | 0.17 | 0.49 |
| 14 | 0.36 | 0.12 | 0.37 | 0.15 |  | 0.45 | 0.14 | 0.29 | 0.12 |
| 15 | 0.13 | 0.49 | 0.24 | 0.14 |  | 0.2 | 0.27 | 0.3 | 0.23 |
| 16 | 0.22 | 0 | 0.13 | 0.65 |  | 0.32 | 0.08 | 0.19 | 0.41 |
| 17 | 0.29 | 0.08 | 0.24 | 0.39 |  | 0.23 | 0.2 | 0.35 | 0.22 |
| 18 | 0.1 | 0.42 | 0.34 | 0.14 |  | 0.23 | 0.27 | 0.36 | 0.14 |
| 19 | 0.16 | 0.01 | 0.09 | 0.74 |  | 0.2 | 0.04 | 0.32 | 0.44 |
| 20 | 0.28 | 0.11 | 0.47 | 0.14 |  | 0.39 | 0.17 | 0.3 | 0.14 |
| 21 | 0.17 | 0.35 | 0.3 | 0.18 |  | 0.26 | 0.25 | 0.24 | 0.25 |

**Table S3.** Synthesized marginal probabilities of library D2 and D3.

| **Library**<br>**D2** | **A** | **T** | **C** | **G** | **Library**<br>**D3** | **A** | **T** | **C** | **G** |
| --- | --- | --- | --- | --- | --- | --- | --- | --- | --- |
| 1 | 0.13 | 0.05 | 0.40 | 0.42 |  | 0.22 | 0.10 | 0.42 | 0.25 |
| 2 | 0.18 | 0.50 | 0.27 | 0.05 |  | 0.22 | 0.29 | 0.34 | 0.15 |
| 3 | 0.20 | 0.21 | 0.27 | 0.32 |  | 0.24 | 0.32 | 0.25 | 0.19 |
| 4 | 0.14 | 0.03 | 0.18 | 0.65 |  | 0.26 | 0.14 | 0.12 | 0.48 |
| 5 | 0.22 | 0.36 | 0.28 | 0.14 |  | 0.19 | 0.39 | 0.25 | 0.17 |
| 6 | 0.27 | 0.27 | 0.24 | 0.22 |  | 0.21 | 0.26 | 0.21 | 0.32 |
| 7 | 0.35 | 0 | 0.14 | 0.51 |  | 0.22 | 0.09 | 0.15 | 0.54 |
| 8 | 0.13 | 0.19 | 0.36 | 0.32 |  | 0.32 | 0.22 | 0.32 | 0.14 |
| 9 | 0.20 | 0.34 | 0.30 | 0.16 |  | 0.24 | 0.20 | 0.25 | 0.31 |
| 10 | 0.13 | 0 | 0.06 | 0.81 |  | 0.25 | 0.08 | 0.41 | 0.26 |
| 11 | 0.27 | 0.14 | 0.21 | 0.38 |  | 0.33 | 0.38 | 0.18 | 0.11 |
| 12 | 0.16 | 0.31 | 0.35 | 0.18 |  | 0.21 | 0.29 | 0.26 | 0.24 |
| 13 | 0.09 | 0 | 0.08 | 0.83 |  | 0.28 | 0.07 | 0.16 | 0.49 |
| 14 | 0.37 | 0.14 | 0.35 | 0.14 |  | 0.45 | 0.16 | 0.27 | 0.12 |
| 15 | 0.12 | 0.53 | 0.22 | 0.13 |  | 0.19 | 0.31 | 0.28 | 0.22 |
| 16 | 0.22 | 0 | 0.12 | 0.66 |  | 0.32 | 0.09 | 0.18 | 0.41 |
| 17 | 0.30 | 0.10 | 0.24 | 0.36 |  | 0.23 | 0.24 | 0.33 | 0.20 |
| 18 | 0.1 | 0.45 | 0.33 | 0.14 |  | 0.22 | 0.30 | 0.35 | 0.13 |
| 19 | 0.16 | 0.01 | 0.09 | 0.74 |  | 0.2 | 0.04 | 0.32 | 0.44 |
| 20 | 0.29 | 0.14 | 0.44 | 0.14 |  | 0.38 | 0.19 | 0.29 | 0.14 |
| 21 | 0.16 | 0.41 | 0.28 | 0.18 |  | 0.25 | 0.30 | 0.23 | 0.22 |
| Total reads assessed by deep sequencing: 193228 (library D2) and 212388 (library D3) |  |  |  |  |  |  |  |  |  |

**Table S4.** Experimental viral titers of glia-infectious AAV variants from ML-D2.

| **Variant** | **Experimental Viral Titer**<br>**(vg/μL)** |
| --- | --- |
| Variant 1 | $2.84 \times 10^{12}$ |
| Variant 2 | $5.97 \times 10^{12}$ |
| Variant 3 | $9.14 \times 10^{12}$ |

**Figure S1:**
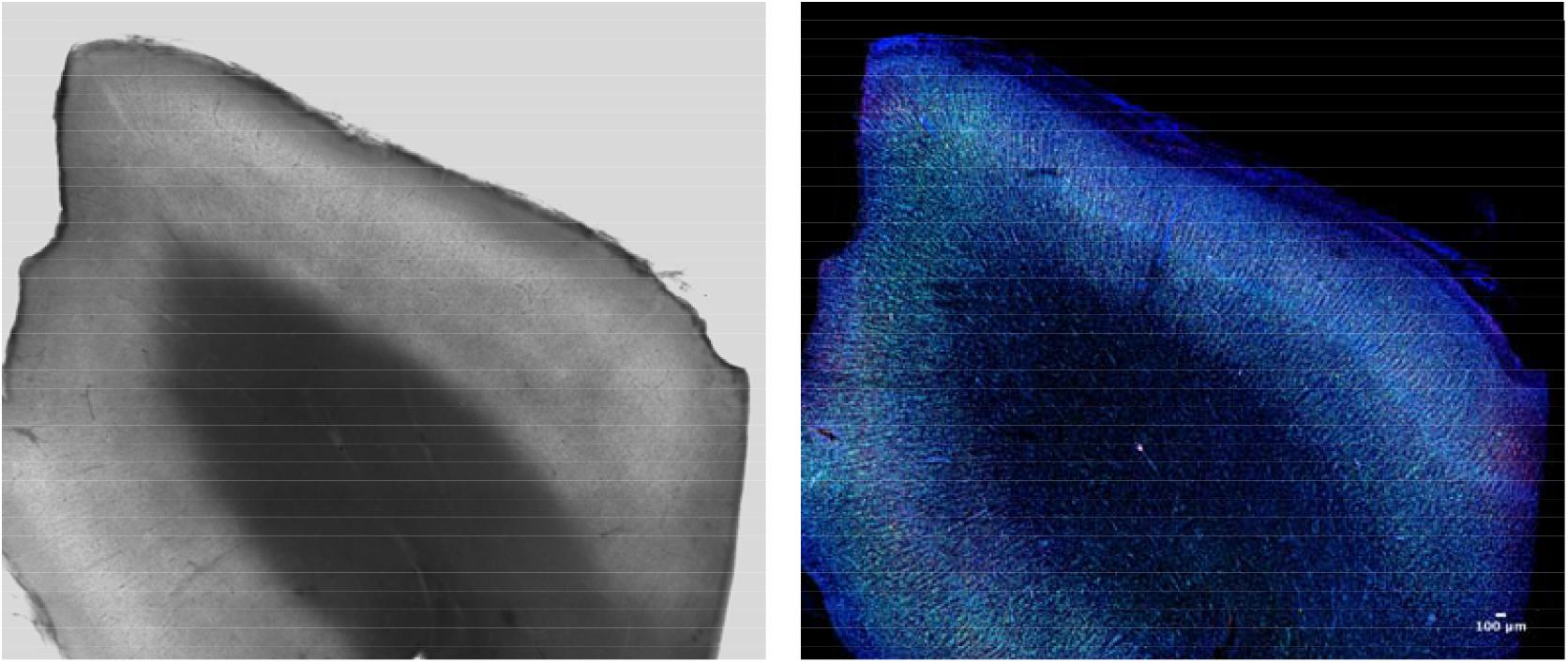
Characterization of primary adult brain section in culture. Brightfield (left); Immunostaining (right): DAPI (blue), Nissl (Cyan), and NeuN (magenta). The middle area is white matter, and the surrounding area is grey matter.

**Figure S2:**
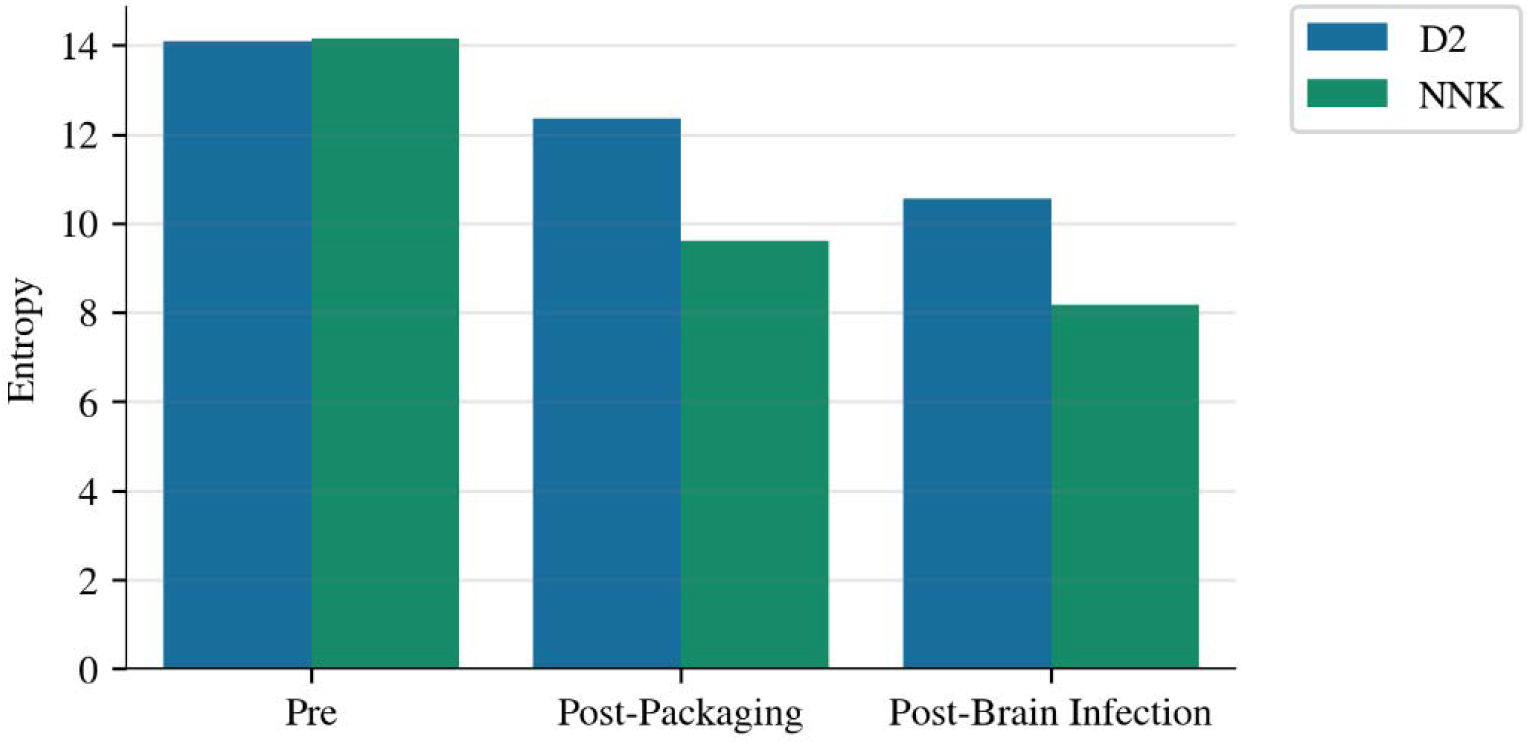
Entropy (diversity) comparison between synthesized NNK and ML-designed D2 libraries after packaging and infection of primary adult brain tissue. D2 presents a comparable level of initial diversity (pre) to that of NNK but outperforms NNK after both packaging (post-packaging) and primary brain infection (post-brain infection).

**Figure S3:**
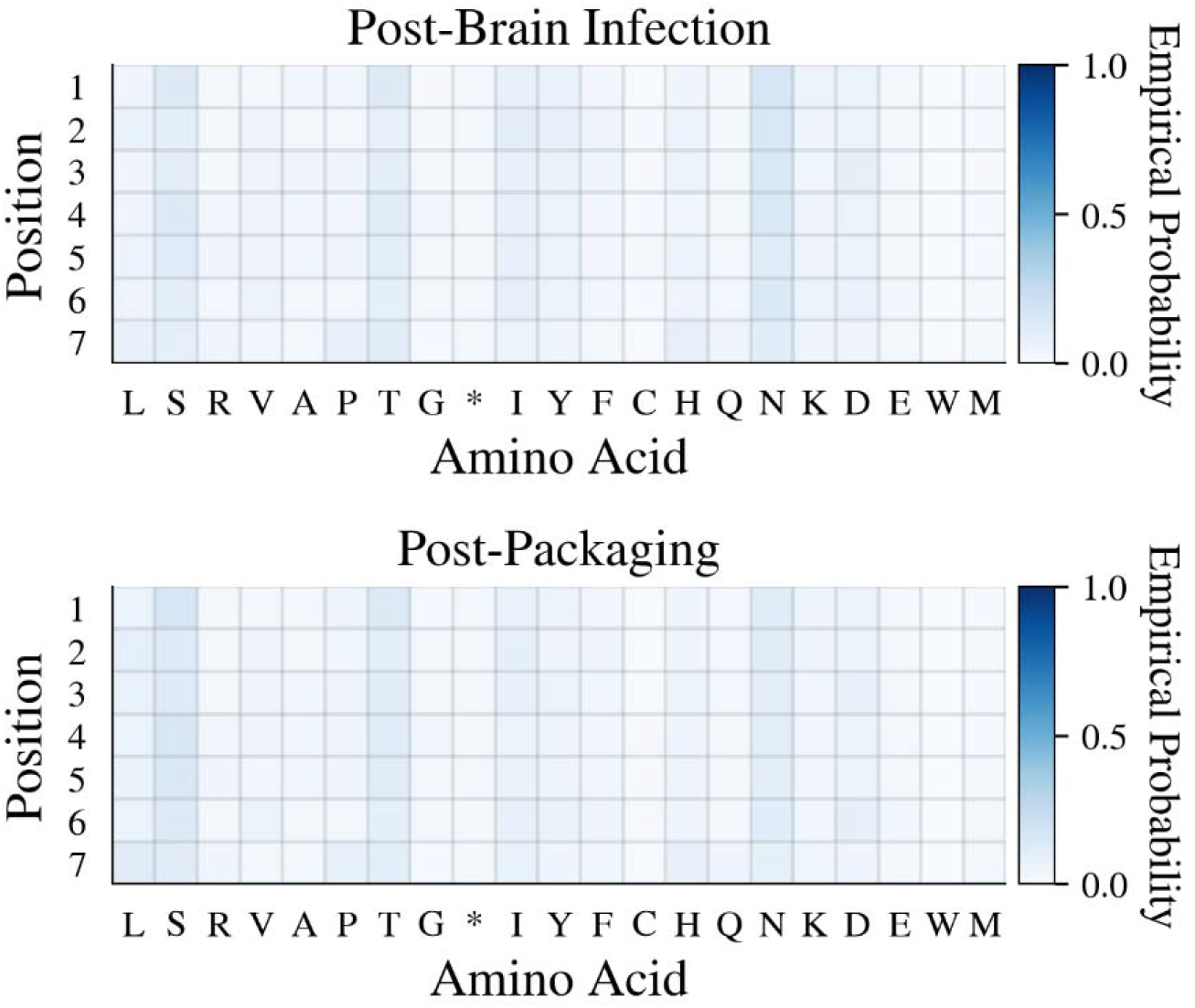
NNK marginal probabilities of amino acids at each position after packaging and primary brain selection.

**Figure S4:**
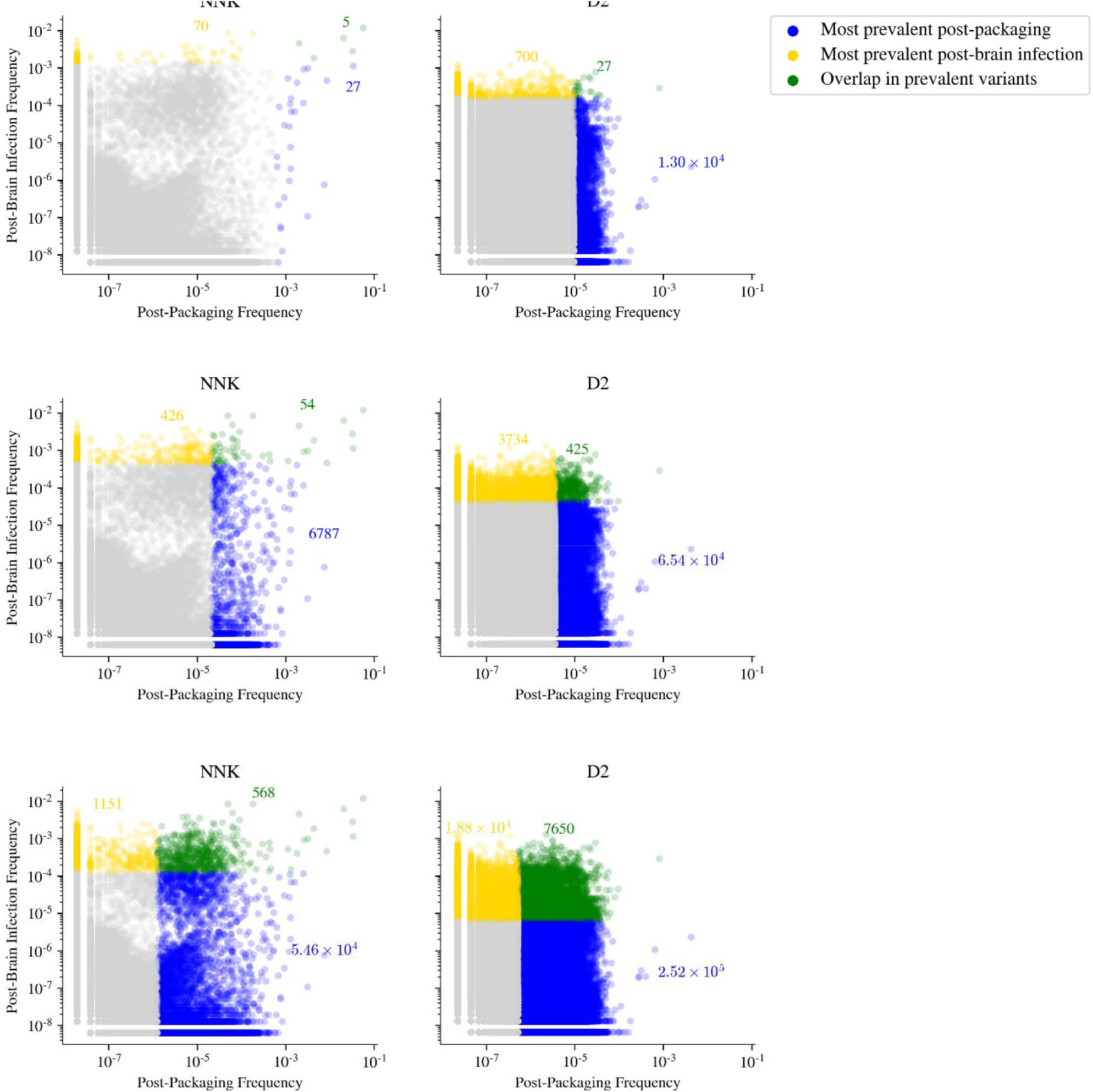
Scatterplots of individual variants’ (log) prevalence after packaging and primary brain selections. From top to bottom: 20%, 50%, and 80% of total reads are assigned colors.

### Supplementary Methods

#### Maximum entropy design of unconstrained libraries

In the main text, we consider constrained library designs, where one specifies experimental control knobs, such as the marginal probabilities of observing each amino acid at each position. Contrasting the constrained libraries, are unconstrained ones, where one constructs a list of oligonucleotide sequences that comprise the library. Unconstrained libraries provide more control over the contents of the library than constrained libraries but are substantially more expensive per oligonucleotide (each of which must be synthesized). Therefore, in considering constrained *vs* unconstrained libraries, one is trading off control for library size. Note that technically, a fully unconstrained library is the probability distribution itself, *p*_*λ*_,(*x*) and that in drawing samples from such a distribution, the resulting library becomes an approximation to the unconstrained library in the sense of having only finitely many samples.

Although we did not experimentally realize any constrained libraries in this work, here we demonstrate that it is possible to apply our maximum entropy formulation to the design of unconstrained libraries. For the purposes of comparison, we exactly compute the entropy and mean predicted log enrichment of the maximum entropy library defined by **Equation 4** by enumerating all possible 7mer insert sequences and evaluating the same predictive model, *f*(*x*), used to design the constrained libraries of Figure 3 for each sequence. In general, even when it is not possible to fully enumerate the relevant sequence space, it is conceptually straightforward to build a list of sequences that approximates the maximum entropy library by sampling from this distribution with, for instance, Markov Chain Monte Carlo (MCMC) algorithms. This resulting set of samples represents a particle-based approximation to Equation 4 and thus will approximately respect the Pareto optimal property of the maximum entropy library.

Figure S6, below, shows the entropy and mean predicted log enrichment for unconstrained libraries corresponding to 404 different settings of *λ*. We can see that unconstrained library construction allows one to build a library with greater diversity at the same level of predicted fitness as constrained libraries. As oligonucleotide synthesis becomes cheaper, unconstrained library synthesis will became correspondingly cheaper. Therefore, our results suggest that at some point, it is likely that unconstrained libraries will become the libraries of choice.

**Figure S5:**
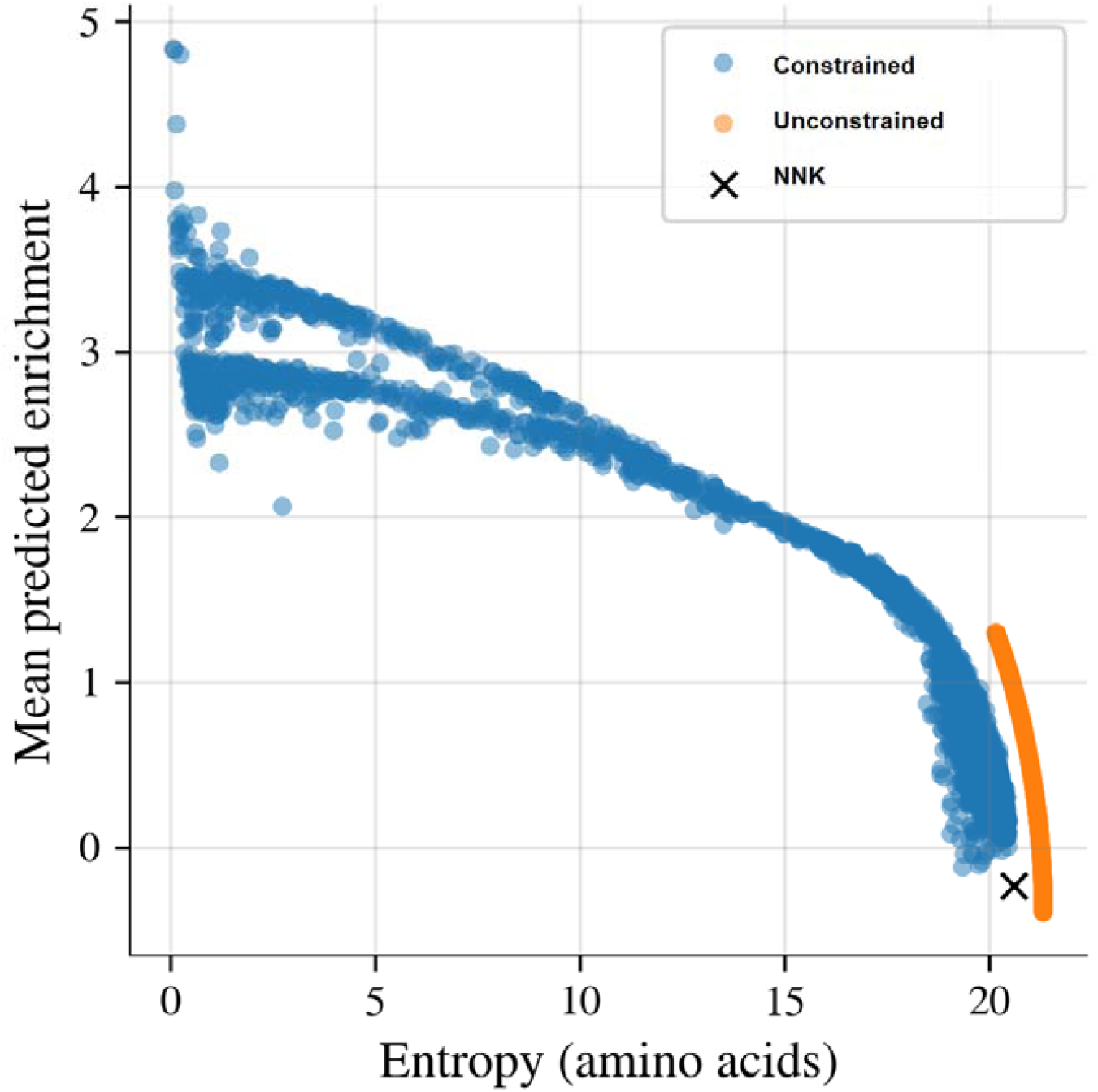
Comparison of maximum entropy unconstrained and constrained libraries. Blue points are identical to the points in Figure 3a, with the colors removed. Orange points represent unconstrained libraries defined by the distribution in Equation 4 with the same values as used to construct the constrained libraries.

#### Gradients for maximum entropy constrained library

To solve the non-convex objective (**Equation 6**) for the library parameters,, we use the Stochastic Gradient Descent (SGD) algorithm, which requires computing the gradient

The gradient of the entropy is given by

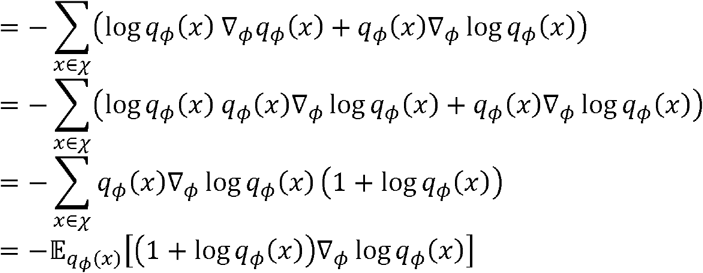

where in the third line we used the equality ∇_*ϕ*_ *q*_*ϕ*_ (*x*) = *q*_*ϕ*_ (*x*) ∇_*ϕ*_ log*q*_*ϕ*_ (*x*). For 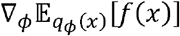, we use the equality 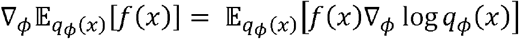 which is well-known from its use in the score function estimator [43] (sometimes also called the ‘log derivative trick’). We then have

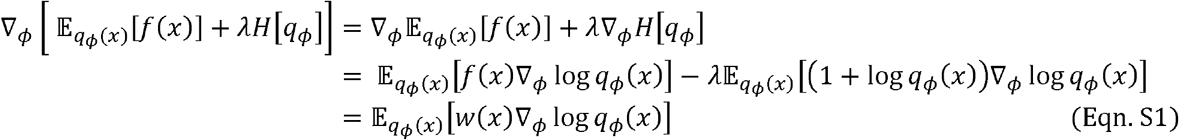

where *w*(*x*) :=*f*(*x*) − *λ*(1 + log*q*_*ϕ*_ (*x*)). The individual components of ∇_*ϕ*_ log*q*_*ϕ*_ (*x*) are given by

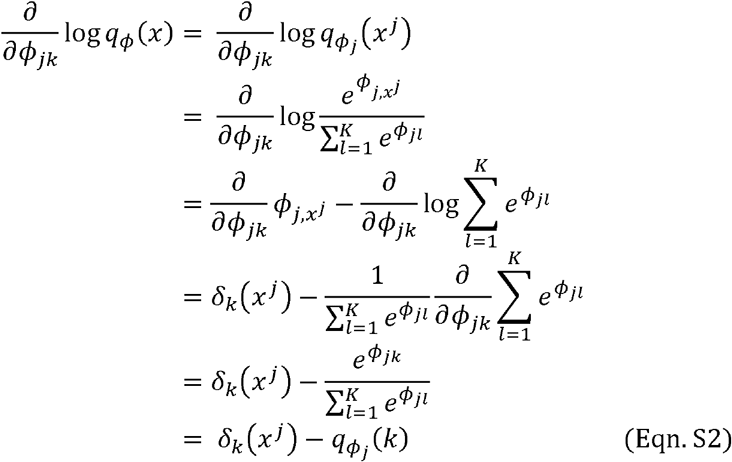

Using **Equation S2** within **Equation S1** gives **Equation 8**.

